# Cataloguing the mouse small intestinal transcriptome in duodenum and ileum during the first postnatal month

**DOI:** 10.1101/2024.09.25.612672

**Authors:** Luiz Fernando Silva Oliveira, Radhika S. Khetani, Nicholas Makogonov, Elizabeth S. Partan, Yu-Syuan Wu, Venkata Siva Dasuri, Amanda W. Harrington, Oluwabunmi Olaloye, Jeffrey Goldsmith, David T. Breault, Liza Konnikova, Shannan J. Ho Sui, Amy E. O’Connell

## Abstract

In the first postnatal month, the developing mouse intestine shifts from an immature to a mature intestine that will sustain the organism throughout the lifespan. Here, we surveyed the mouse intestine in C57Bl/6 mice by bulk RNA-Seq to evaluate the changes in gene expression over time from the day of birth through 1 month of age in both the duodenum and ileum. Using trends identified in the RNA-Seq analyses, we further evaluated expression of epithelial and mesenchymal cell markers, epithelial regulators, and immune cell markers. We confirmed key changes with qRT-PCR and immunofluorescence. In addition, we compared some findings to humans using human intestinal biopsies and organoids. This dataset can serve as a reference for other groups considering the role of single molecules or molecular families in early intestinal and postnatal development, expanding the limited literature on postnatal gene expression in the developing intestine.

## Introduction

The post-embryonic, developing intestine remains a dynamic organ that undergoes important molecular shifts as the crypts develop and the cellular composition matures. Exposures to the microbiome and nutrients cause additional programmatic changes. In the mouse, the postnatal period between day 0 and week 4 approximates the developmental sequence of premature human infants around 28-34 weeks’ gestation, as both species experience additional crypt-villous maturation and the development of Paneth cells during these intervals (*1–5*). While many groups have described the developmental sequence of gastrulation (*6–10*) and gut tube formation (*11, 12*), as well as embryonic generation of the foregut (*13*), midgut (*13–15*), and hindgut (*16*), less is known about the second half of pregnancy in humans or the postnatal developmental sequence in either species.

Studying human development during the second half of pregnancy has been challenging, owing to the difficulty of obtaining tissue from fetal intestine after about 24 weeks. Even when tissue can be obtained from premature infants, it is requisite that a disease is present, causing the need for surgical intervention – whether that be a congenital malformation of the intestine (duodenal atresia, gastroschisis, etc.) or an acute disease process (spontaneous perforation, necrotizing enterocolitis). Mouse models are frequently used to help us understand human intestinal development, however, most studies of intestinal development in mice focus on the in-utero period and important questions remain about postnatal development (*17*). In general, existing postnatal studies have focused on the role of single genes in modulating postnatal development (*18–20*). While there is a rich body of literature examining the impacts of the microbiome on postnatal intestinal development (*18, 21, 22*), comprehensive analyses of the postnatal intestinal sequence are lacking, especially compared to the breadth of studies on prenatal development. A study in Balb/c mice showed that cell proliferation is robust in the neonatal mouse and exceeds cell loss until around week three, when more adult-like homeostatic kinetics are attained (*23*). Another study in B6 mice looked at transcript patterns in crypts versus villi in the postnatal period using RNA transcripts from the medial small intestinal segment (*24*).

We are interested in understanding fundamental biological processes that occur during intestinal development in the neonatal period. We performed a time course analysis of postnatal development using bulk RNA-Seq of the full thickness mouse intestine, which includes epithelium and mesenchymal tissues. We also examined differences in gene expression patterns of the duodenum versus ileum. Findings were confirmed using other modalities and human tissue biopsies were used for comparison in select instances.

## Results

### Experimental approach

For this study we used mouse intestinal tissue from the most proximal 5cm (duodenum) and most distal 5cm (ileum) small intestine to generate bulk RNA for RNASeq (Figure 1A.) We used similar but unique samples for qRT-PCR confirmation of select results. We also used histologic samples obtained from full length small intestinal Swiss-rolls from matching-aged mice for immunofluorescence analysis. We oriented these rolls in the same fashion and confirmed the appropriate location in the ileum both by orientation on the slide and by geographic features of the ileum (shorter villi, increased goblet cells, Peyer’s patches). For human assays, since we did not have fresh tissue for RNA available, we used small intestinal histology sections obtained from premature-born infants for RNAScope and immunofluorescence. To do bulk RNA analyses, we used epithelial-only organoids obtained from premature-born infant within the 72 hours of life at the specified gestational age.

**Figure 1.**
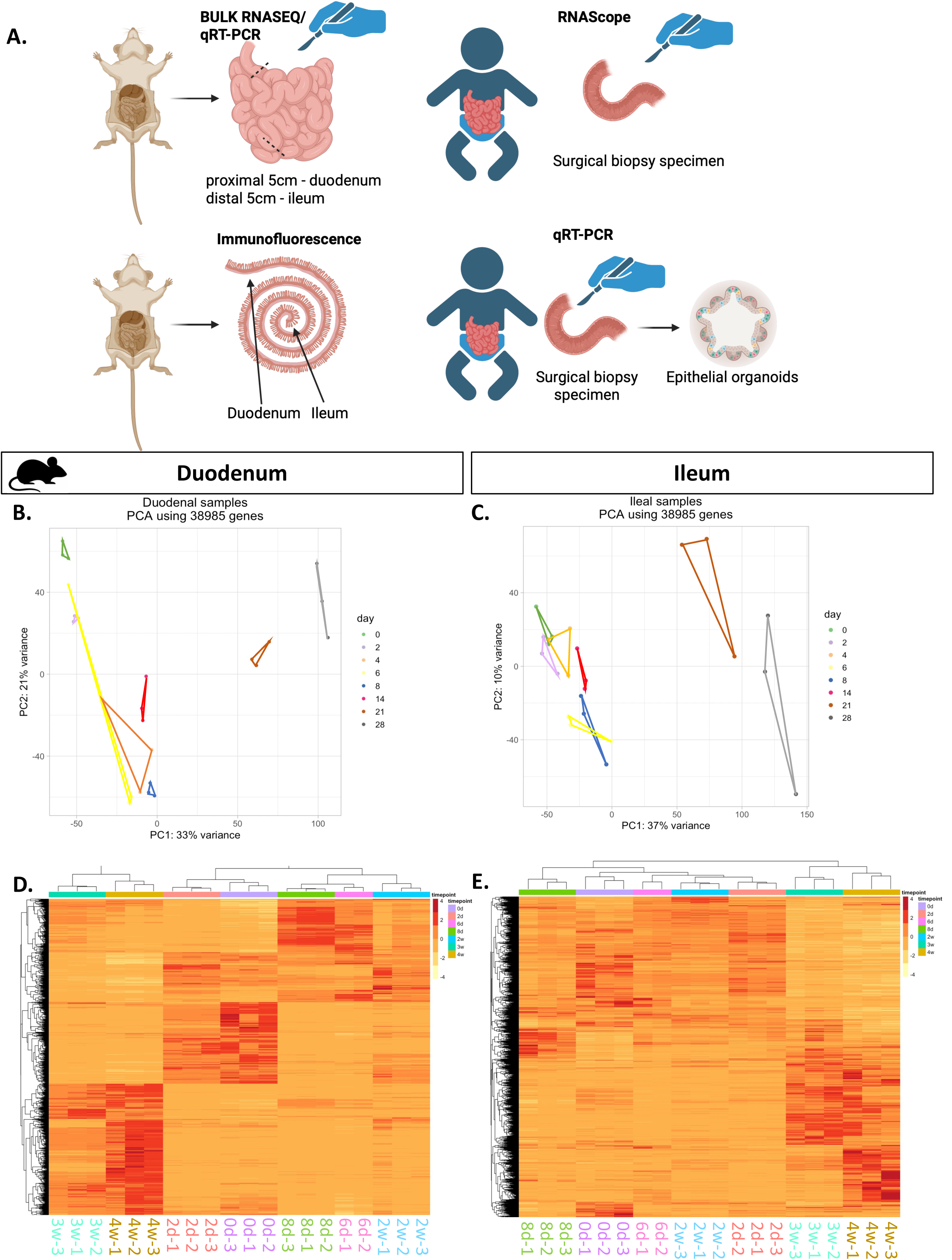
Sample origins, PCA and hierarchical clustering. (A) Schematic of sample collection for mouse and human samples. B & C Principal component analysis of all Duodenum samples (B) and all Ileum samples (C). D. & D. Heat map of differentially expressed genes displayed with hierarchical clustering in all Duodenum samples (D) and in all Ileum samples (E). DE genes (time course using LRT) were defined as adjusted P-value <= 0.005.

### The quality of our mouse small intestinal RNA-Seq dataset was high, and samples clustered together by developmental timepoint

To examine the quality of the RNA-Seq dataset, we examined the total reads per sample; read numbers ranged from 56 to 185 million reads per sample (Supplemental Fig. 1A). In general, ileal samples had fewer reads than duodenal ones. We also evaluated the exonic mapping rate, and most of the samples met our target threshold of 75%, although four of the ileal samples had mapping rates slightly below 70% (Supp. Fig. 1B). Principal component analysis (PCA) of all samples showed that duodenal samples and ileal samples intermixed and did not segregate by tissue type in early development but segregated more at later time points (Supp. Fig. 1C), suggesting that these regions were similar in gene expression through the first two weeks of life.

To analyze the clustering within each group, we also performed PCA for samples from each tissue independently. Duodenal samples generally clustered well by time point and followed a temporal sequence in the principal driver of variance, PC1 (Supp. Fig. 1D). The samples from the ileum similarly clustered well and followed a temporal pattern in PC1 (Supp. Fig. 1D). The 2-day through 2-week samples were more closely related to each other. In the ileum, the 3– and 4-week samples again clustered apart from the rest of the group, but the 0d samples clustered similarly to the 2-day through 2-week samples. However, the samples from day 4 did not cluster tightly, particularly in the duodenum, and there was an outlier in the day 6 samples that did not cluster with the other two in duodenum or ileum. Given that the younger timepoints generally clustered tightly, and we had many time points, we opted to drop the day 4 samples and one of the day 6 samples from the rest of our analyses, to focus on the time points that gave us good replication between the samples and had higher reliability.

Analysis of gene expression using significantly differentially expressed (DE) genes (adjusted p-value/q-value <= 0.005) demonstrated that significantly different gene expression in the duodenum was similar in 0-day and 2-day mice, 6-day, 8-day and 2-week mice, and 3-week and 4-week mice, as displayed in heatmap format (Fig. 1D). The similarities between DE gene expression in the ileum were less linear with 0-day and 8-day aligning independently and 2-day, 6-day and 2-week samples appearing more similar in the DE gene profile (Fig. 1E). 3-week and 4-week profiles were also similar, although 4-week samples appeared to have an additional cluster of highly DE genes compared to the 3-week samples (bottom right). See Supplemental File 1 for gene counts and differentially expressed gene lists.

### Gene expression trends in the duodenum and ileum highlight key developmental gene expression changes

We next evaluated all differentially expressed genes across time in the duodenum (Supp. Fig. 2; Supp. File 2) and ileum (Supp. Fig. 3; Supp. File 3) and asked whether there were patterns in gene expression trends that were unique in each dataset. Using the top 3000 differentially expressed genes in each tissue (adjusted P-value threshold used <= 0.005), we identified 20 gene sets with correlated gene expression trends. We then called out gene set clusters in the ileum that either were low in the pre-weaning postnatal period (day 0-day 14) and then increased at week 3-4, or that were elevated in earlier times points and then decreased significantly as the intestine matured. We performed gene ontology (GO) enrichment analyses on these smaller genes sets (Fig. 2) to identify overrepresented GO terms. Notably, genes for immunity were predominant in one of the clusters (cluster 3). Other clusters that increased in the post-weaning period encoded genes for metabolism and intracellular trafficking (cluster 2) and ATP/energy use (cluster 4). Gene groups that were elevated prior to weaning included tissue development (angiogenesis, metabolism, catabolism genes) (clusters 1), cell migration genes (cluster 11), DNA repair (cluster 7), and vacuole development (cluster 5). One gene set, cluster 17, was low initially at day 0 and day 2, then increased from day 6 to 2 weeks, then decreased again in weeks 3 and 4, and this was enriched for genes associated with tissue development including extracellular matrix and smooth muscle gene families.

**Figure 2.**
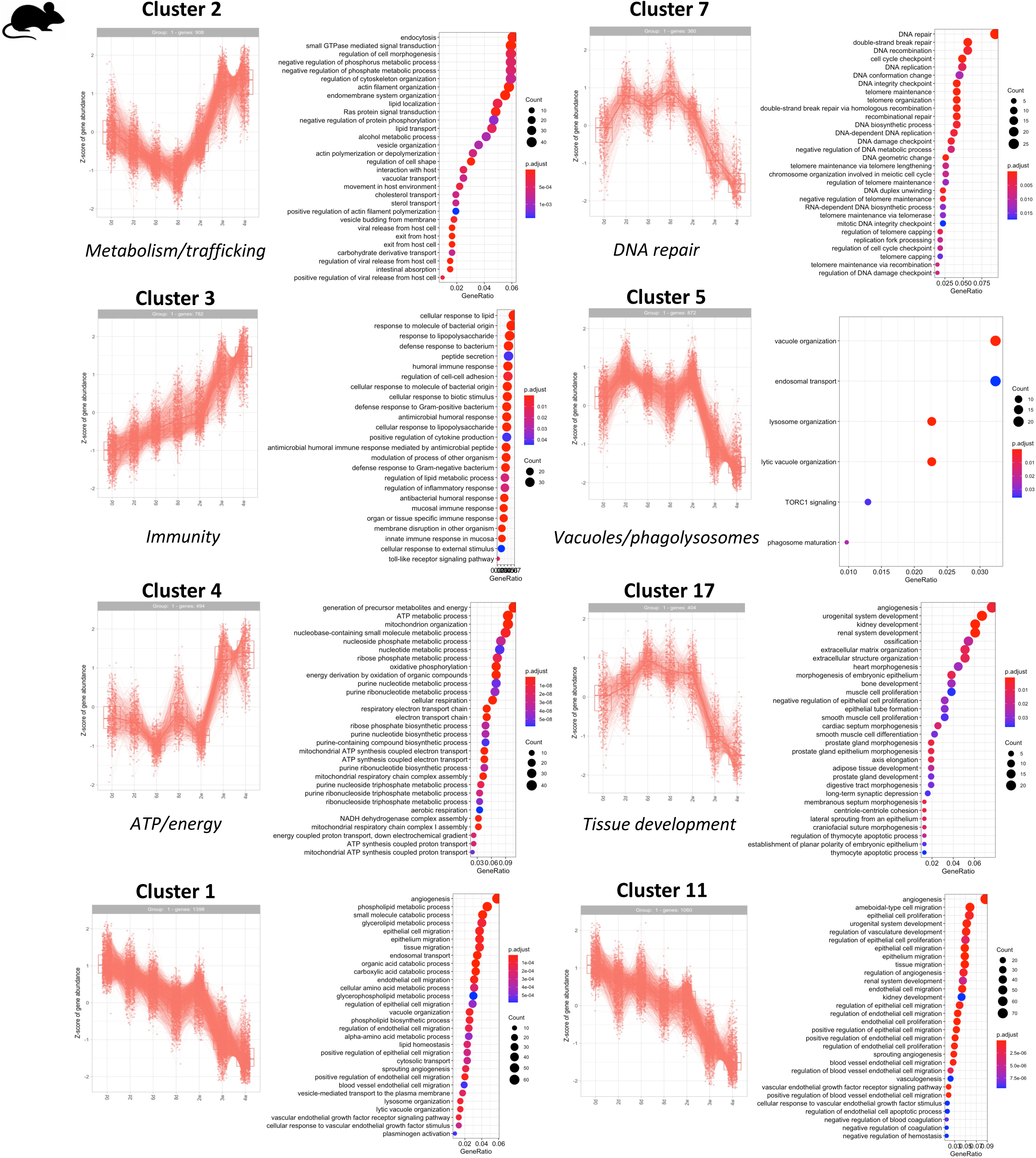
Gene ontology enrichment analysis of selected gene expression clusters. For each gene cluster, the expression pattern, number of genes present, and abundance score are indicated on the left. On the right, the top (top 30, except for cluster 5) overrepresented gene ontology terms in the cluster are indicated. Gene ratio on the y-axis indicates the proportion of your significantly altered genes that are involved in a specific biological pathway or function.

**Figure 3.**
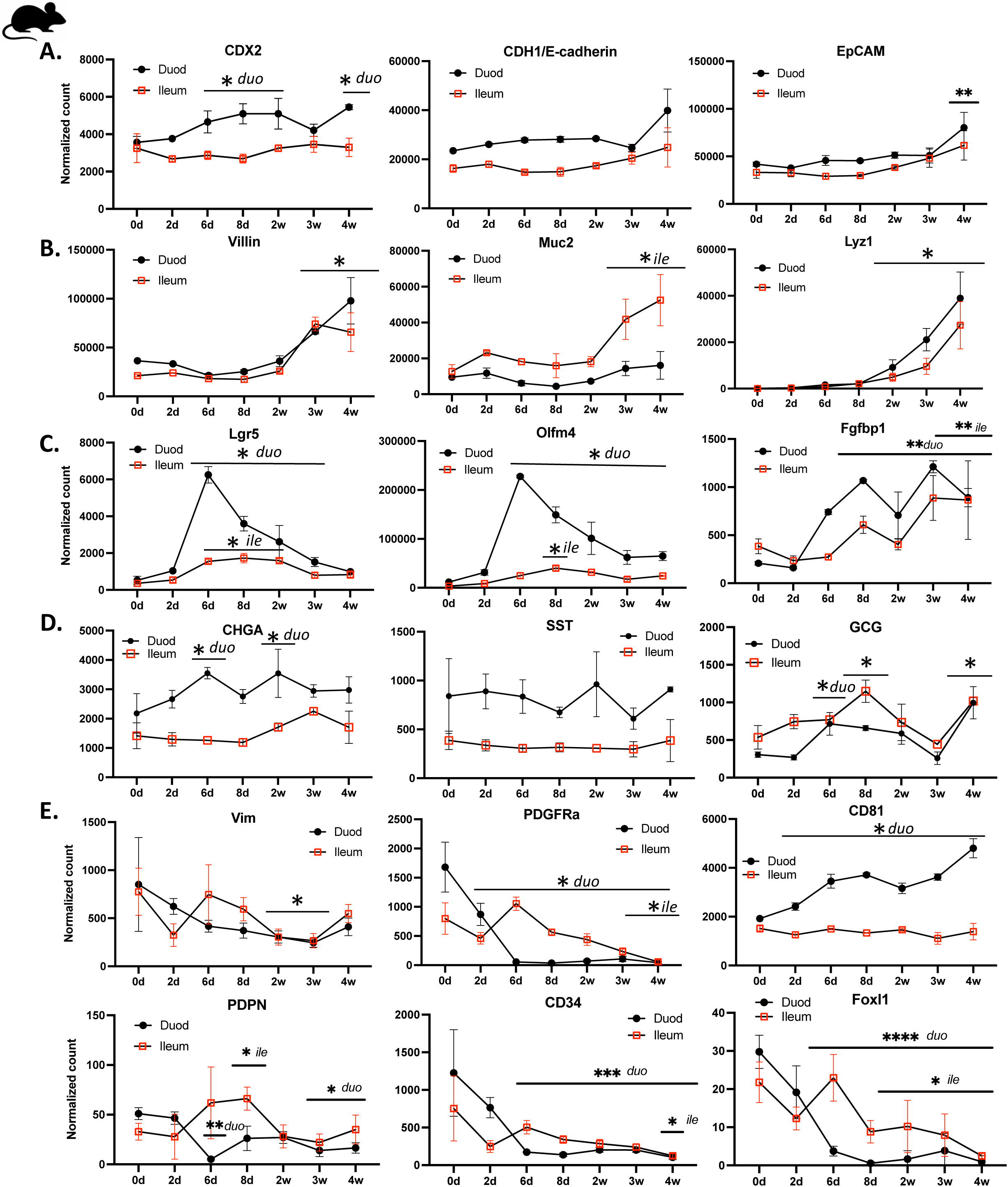
Expression of intestinal epithelial and mesenchymal markers. Gene expression in the duodenum (black) and ileum (red) by normalized gene count across the first month of postnatal life. Highlighted genes include general markers of the intestine (A), intestinal epithelium and secretory populations (B), intestinal stem cells (C), enteroendocrine cells (D), and intestinal mesenchyme (E). Statistical analysis was by 2-way ANOVA compared to day 0. (F) qRT-PCR analysis of select genes from duplicate intestinal samples at the indicated time points. *p< 0.05, *duo – duodenum significant only, *ile-ileum significant only, ***p<0.001.

### General intestinal markers are stable throughout time points

Given that we used full-thickness biopsies of the intestinal tissue for our analyses, we wanted to ensure that tissue samples were consistent throughout our dataset as an internal validation. We plotted expression of intestine specific markers *CDX2* (specification marker), *CDH1* (E-cadherin, epithelial marker) and *EPCAM* (epithelial cell adhesion molecule, epithelial marker) and saw that expression for duodenum and ileum samples was relatively stable over time, while both were increased in duodenum versus ileum (Fig. 3A). The increased villus length in the duodenum may explain this relative increase. The overall consistency of expression of these ubiquitous markers helps to establish reliable tissue sampling between time points.

### Epithelial differentiation markers increase over time, while stem cell markers peak around postnatal week 1

One of the genes present in gene cluster 2 was villin, an epithelial marker, suggesting that epithelial villus growth, potentially in the form of increased length or increased density, is represented by cluster 2. To examine this in more detail, we called out specific markers of epithelial cell sub-types. Expression of epithelial differentiation markers *Vil1* (Villin, all epithelium) and *Muc2* (Mucin 2, goblet cells) remained relatively low over the first 2 weeks of life, then increased significantly in weeks 3 and 4 (Fig. 3B). Similarly, and as described previously (*1–3, 5*), expression of *Lyz1* (Lysozyme 1, Paneth cells) also remained low in the first few weeks of life but was statistically increased from week 2 onward. In contrast to the differentiation markers, conventional intestinal stem cell (ISC) markers *Lgr5*, *Olfm4*, and *Ascl2* peak around 6-8 postnatal days (Fig. 3C). Meanwhile, expression of markers for enteroendocrine cells, *Chga* (chromogranin A), *Sst* (somatostatin), and *Gcg* (glucagon), did not follow a consistent pattern (Fig. 3D). It is important to note, however, that duodenal and ileal samples were processed in separate analyses and, for this figure, levels of gene expression may not be completely relative. Instead, the goal of graphing together is to compare expression trends between the two issues. Further, these markers are highly associated with these cell subtypes, however they may be expressed in smaller amounts in other lineages, so the interpretation of populations may be confounded.

### Mesenchymal markers also demonstrate varied expression by time point

Cluster 11, which had genes that were higher from P6-P14, included *Pdgfra*, a marker for key mesenchymal cell populations. *Pdgfra* (platelet-derived growth factor A), is expressed in several intestinal mesenchymal cell types as well as vascular cells, and expression peaked at day 0 in the duodenum and then fell off rapidly (Fig. 3E). In the ileum, in contrast, expression was stable until 3 weeks. Expression of *Vim* (vimentin), a broad marker for the intestinal mesenchyme, was highest at day 0 and then significantly decreased from day 0 by 2 weeks in the ileum and 3 weeks in the duodenum, then stayed lower. *Cd81*, a marker for stromal telocytes (*25–27*), rose over time in the duodenum but remained stable in the ileum. These findings suggest a shift in the postnatal fibroblast population in the first postnatal month.

### Confirmation of key findings by qRT-PCR

To confirm reproducibility of our results across samples and techniques, we collected samples from other mice at select time points and measured RNA of several of the genes of interest using qRT-PCR (Fig. 3F). Indeed, expression of *Olfm4*, *Muc2*, and *Pdgfra* correlated with the gene expression data observed in the bulk RNASeq analysis. Olfm4 expression was highest at day 6, although by qRT-PCR duodenum and ileum levels were similar. *Muc2* expression was higher in the ileum throughout and significantly higher at week 3, consistent with RNASeq. *Pdgfra* expression decreased at the later time points.

### Comparison of data to published subpopulation datasets

We next used published mouse intestine single cell RNASeq datasets for epithelial (*28*) sub-populations and mesenchymal (*29*) subpopulations to perform cellular deconvolution of our data. This enables us to use gene sets that are known to designate particular cellular subpopulations to approximate the presence of subpopulations using our bulk RNASeq data (*30*). Proximal and distal enterocytes increased over time in duodenum and ileum, consistent with villus lengthening (Fig. 4A). Enteroendocrine cell expression was relatively stable in the ileum but increased at day 6 in the duodenum. Goblet and tuft cells were overall stably expressed in both regions, and Paneth cells, in alignment with previous studies (*31*), increased with maturity.

**Figure 4.**
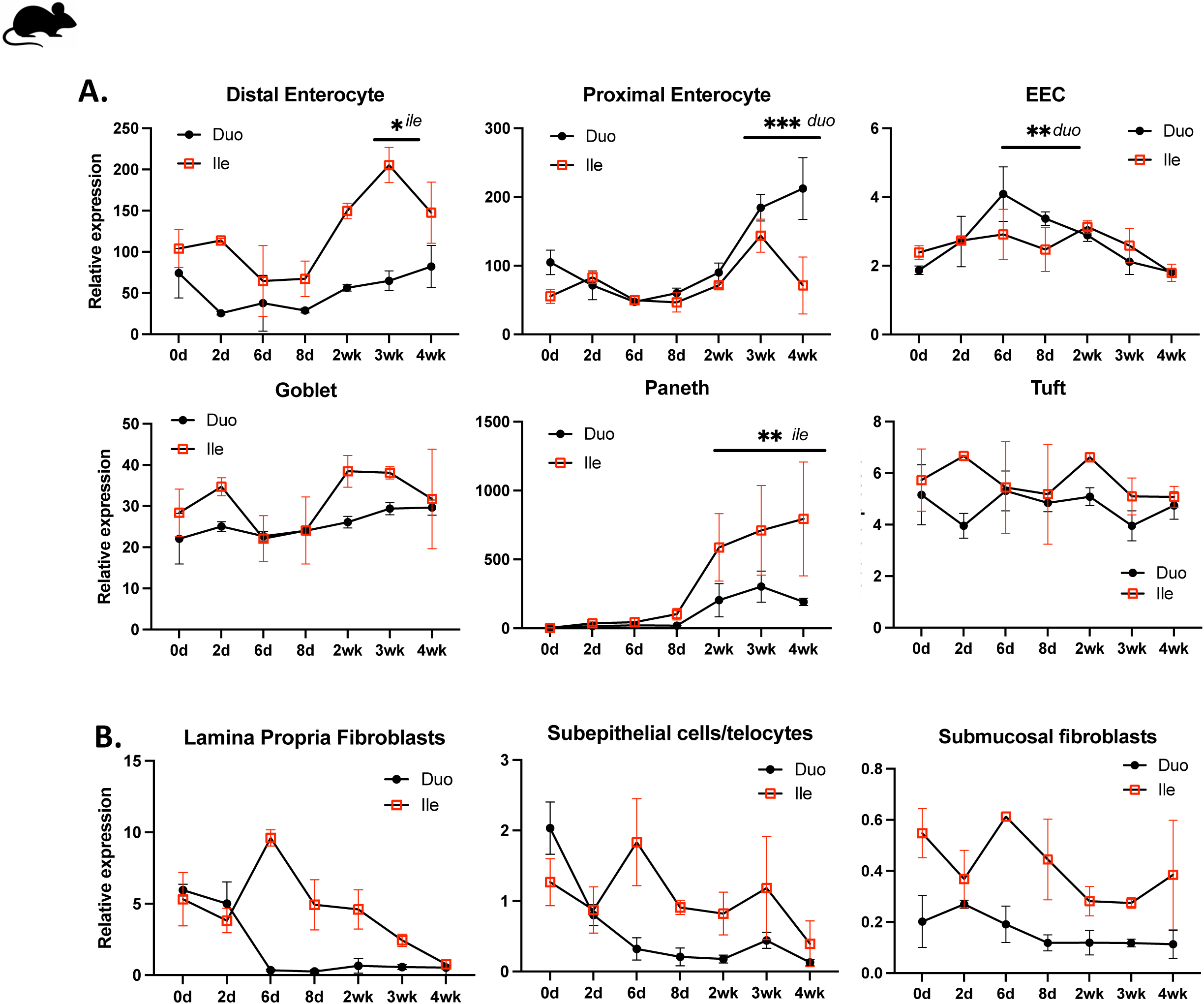
Deconvoluted datasets for key epithelial and mesenchymal subsets. (A) Epithelial cell subpopulations using published data for key subpopulation markers. (B) Mesenchymal subpopulations using published data for early life subepithelial fibroblast subpopulations.

The ileum had a higher overall composition of fibroblast subtypes than the ileum (Fig. 4B). Lamina propria fibroblasts decreased over time in the duodenum, but initially increased in the ileum at day six before decreasing. Subepithelial telocytes also fell steadily in the duodenum and with higher variation over time in the ileum. Submucosal fibroblasts were more consistent over time on both segments.

### Expression of stem cell markers in the developing human intestine

We next sought to compare expression of select epithelial and trophic factors in intestinal epithelium from premature-born infants. With the caveat that samples can only be obtained in the instances of a clinical need for surgery, we used biopsy samples from intact, healthy-appearing tissue from infants who were diagnosed with NEC and had tissue collected from the ileum or jejunum (Table 2). In a 26-week infant, *LGR5* expression was already well organized at the crypt base, highlighting the ISCs (Fig. 5A). Expression was still limited to the crypt at 29 weeks but was more widespread at 35 weeks.

**Figure 5.**
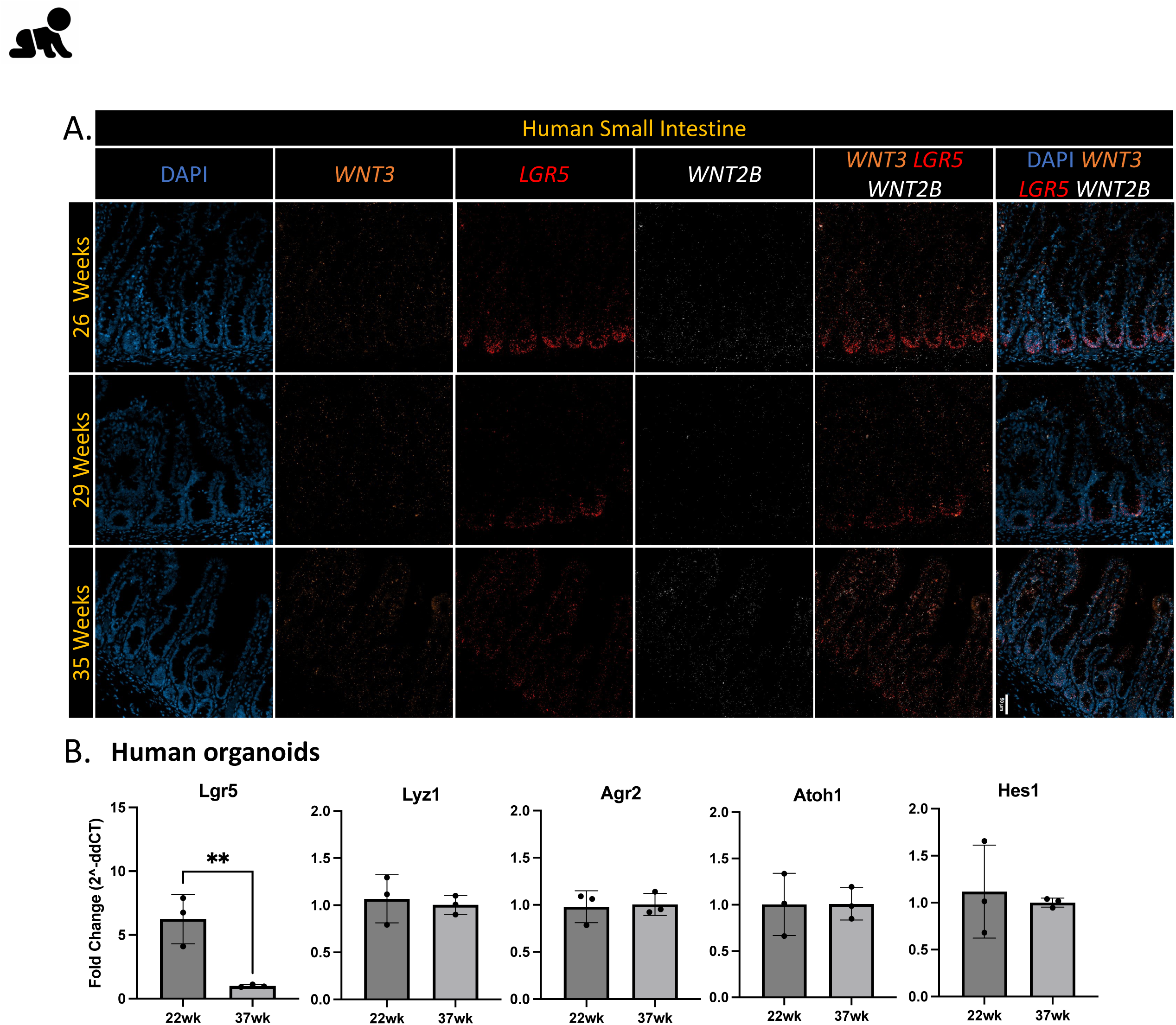
Expression of key epithelial genes in human tissues. (A) RNAScope of biopsied human tissue from the indicated gestational ages (Clinical data from the specimens is indicated in Table 2). DAPI is in blue, *LGR5* in red. (B) RNA expression in human intestinal organoids from 22-week fetus and 37-week infant by qRT-PCR. ** p<0.01.

**Table 1.** Average Expression Duodenum and Ileum Defensins by RNASeq.

| **DUO-**<br>**DENUM** | **0d** | **2d** | **6d** | **8d** | **2w** | **3w** | **4w** |
| --- | --- | --- | --- | --- | --- | --- | --- |
| Defa24 | 3161.68 | 5729.47 | 11143.71 | 8650.94 | 19732.65 | 27531.96 | 29333.56 |
| Defa23 | 11.26 | 28.66 | 676.38 | 1039.39 | 7571.67 | 16412.25 | 19091.25 |
| Defa3 | 1144.91 | 1730.69 | 5660.92 | 5158.98 | 10520.79 | 14829.51 | 15179.76 |
| Defa30 | 12.65 | 36.65 | 510.36 | 580.77 | 5075.75 | 10160.41 | 10496.18 |
| Defa43 | 12.24 | 26.17 | 283.37 | 480.23 | 1670.80 | 3877.55 | 9970.95 |
| Defa17 | 633.59 | 1065.88 | 3147.14 | 2813.13 | 5475.09 | 7859.05 | 9002.39 |
| Defa36 | 5.85 | 68.65 | 1147.63 | 1186.55 | 6192.45 | 9641.40 | 8622.08 |
| Defa39 | 8.31 | 7.25 | 105.64 | 200.19 | 814.46 | 2727.75 | 7698.69 |
| Defa2 | 1.29 | 5.57 | 332.01 | 506.68 | 4450.94 | 7113.18 | 4614.90 |
| Defa20 | 4.24 | 10.18 | 296.89 | 477.35 | 3271.28 | 5289.79 | 3345.31 |
| Defa38 | 0.00 | 0.00 | 3.26 | 6.38 | 135.53 | 1180.30 | 3158.90 |
| Defa29 | 0.94 | 0.53 | 5.15 | 14.36 | 154.17 | 843.31 | 2753.19 |
| Defa22 | 0.24 | 0.37 | 77.69 | 119.86 | 1334.68 | 2236.71 | 1270.90 |
| Defa21 | 0.00 | 1.51 | 84.71 | 90.17 | 1114.43 | 2070.63 | 1063.54 |
| Defa41 | 0.00 | 2.47 | 76.36 | 106.98 | 863.98 | 1525.23 | 915.75 |
| Defa34 | 0.00 | 5.22 | 89.08 | 149.24 | 715.63 | 1128.59 | 871.47 |
| Defa33 | 0.82 | 0.00 | 52.46 | 140.07 | 927.83 | 1448.12 | 677.01 |
| Defa26 | 6.79 | 9.15 | 48.37 | 68.05 | 248.21 | 414.86 | 406.72 |
| Defa40 | 0.24 | 0.00 | 5.15 | 5.36 | 57.95 | 193.04 | 239.62 |
| Defa37 | 6.01 | 0.00 | 0.00 | 8.25 | 61.89 | 11.92 | 204.96 |
| Defa32 | 0.47 | 0.00 | 27.68 | 53.69 | 404.38 | 537.29 | 178.42 |
| Defa35 | 0.00 | 0.53 | 22.97 | 27.99 | 163.43 | 323.73 | 141.22 |
| Defa28 | 2.10 | 0.79 | 16.41 | 21.86 | 62.19 | 122.87 | 60.84 |
| Defa27 | 0.58 | 0.27 | 3.27 | 7.73 | 31.09 | 69.55 | 18.93 |

| ILEUM | 0d | 2d | 6d | 8d | 2w | 3w | 4w |
| --- | --- | --- | --- | --- | --- | --- | --- |
| Defa2 | 1.26 | 20.91 | 797.78 | 1628.66 | 4994.25 | 6929.98 | 10692.13 |
| Defa23 | 7.23 | 22.66 | 406.95 | 1231.80 | 3115.56 | 5161.23 | 10614.92 |
| Defa20 | 3.44 | 20.28 | 918.37 | 1807.18 | 4878.48 | 7165.43 | 10032.43 |
| Defa43 | 8.01 | 58.53 | 520.80 | 1305.76 | 2210.36 | 3669.14 | 8972.27 |
| Defa36 | 0.00 | 91.48 | 1137.85 | 2125.92 | 4625.80 | 6079.72 | 8914.86 |
| Defa3 | 779.31 | 1554.81 | 2371.39 | 4535.10 | 4800.40 | 6262.30 | 8505.39 |
| Defa30 | 5.20 | 19.81 | 323.37 | 823.52 | 2523.17 | 3758.30 | 8037.65 |
| Defa22 | 0.42 | 1.63 | 270.83 | 545.35 | 2359.44 | 4053.24 | 6277.05 |
| Defa33 | 0.00 | 4.27 | 382.18 | 889.65 | 2527.05 | 4249.49 | 6203.35 |
| Defa17 | 460.76 | 1120.92 | 1352.89 | 2917.78 | 2228.19 | 2922.15 | 5746.62 |
| Defa32 | 0.70 | 18.85 | 350.16 | 852.37 | 2262.58 | 3947.68 | 5029.42 |
| Defa21 | 0.00 | 1.65 | 222.50 | 440.08 | 1841.28 | 3385.60 | 4695.00 |
| Defa5 | 0.00 | 1.10 | 165.88 | 399.26 | 1079.18 | 1687.81 | 2597.65 |
| Defa39 | 1.75 | 4.57 | 54.42 | 200.50 | 491.46 | 1060.32 | 2416.73 |
| Defa29 | 0.42 | 0.67 | 8.08 | 48.50 | 164.22 | 593.30 | 2318.58 |
| Defa34 | 0.79 | 7.62 | 146.94 | 379.82 | 671.08 | 910.92 | 1641.24 |
| Defa41 | 0.21 | 2.20 | 144.78 | 328.82 | 912.89 | 1314.96 | 1575.60 |
| Defa38 | 0.00 | 0.69 | 1.69 | 16.20 | 125.84 | 498.79 | 1476.09 |
| Defa37 | 2.13 | 0.00 | 0.00 | 53.78 | 83.12 | 143.86 | 375.39 |
| Defa35 | 0.33 | 1.29 | 35.27 | 73.63 | 163.27 | 266.81 | 320.28 |
| Defa26 | 3.25 | 7.91 | 30.32 | 84.01 | 119.01 | 158.63 | 275.13 |
| Defa40 | 0.00 | 0.34 | 3.54 | 16.55 | 43.20 | 90.21 | 148.83 |
| Defa28 | 0.87 | 0.34 | 5.66 | 35.06 | 42.54 | 74.22 | 79.85 |
| Defa27 | 0.00 | 0.28 | 2.26 | 10.23 | 17.83 | 32.80 | 35.90 |

**Table 2.**
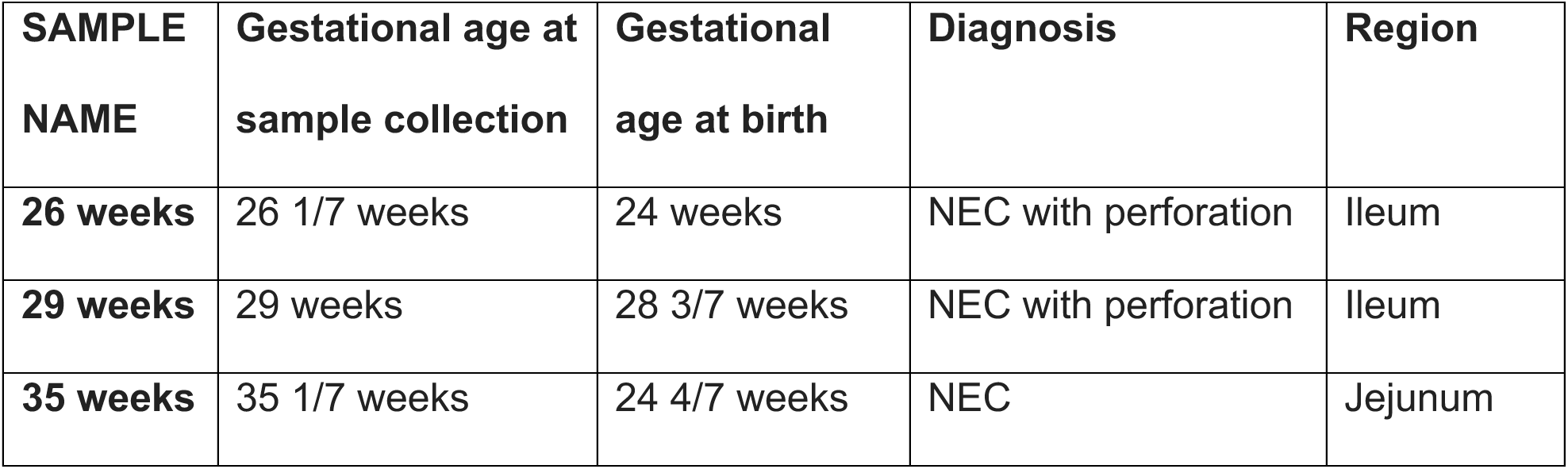
Demographic data from human biopsy samples.

| **SAMPLE NAME** | **Gestational age at sample collection** | **Gestational age at birth** | **Diagnosis** | **Region** |
| --- | --- | --- | --- | --- |
| **26 weeks** | 26 1/7 weeks | 24 weeks | NEC with perforation | Ileum |
| **29 weeks** | 29 weeks | 28 3/7 weeks | NEC with perforation | Ileum |
| **35 weeks** | 35 1/7 weeks | 24 4/7 weeks | NEC | Jejunum |

We similarly generated human epithelial organoids from human intestinal biopsies from a 22-week fetus and a 37-week newborn (Table 3) and evaluated the expression of intestinal lineage markers (Fig. 5B). *LGR5*, the prototypical stem cell marker, was significantly lower in the 37-week organoids, however, *OLFM4* expression was higher in the 37-week organoids compared to the 22-week (2^nd^ trimester) ones. It was unexpected that these intestinal stem cell markers did not correlate. Other lineage markers (*LYZ, AGR2, ATOH1*) were similar between the groups (not shown), however we did not use differentiation media so it is expected that the culture medium would favor maintenance of stem cells over epithelial differentiation.

**Table 3.** Demographic data from human organoids.

| **SAMPLE NAME** | **Gestational age at sample collection** | **Gestational age at birth** | **Gender** |
| --- | --- | --- | --- |
| **22 weeks** | 22 weeks | N/A | Unknown |
| **37 weeks** | 37 2/7 weeks | 37 5/7 weeks | Female |
| **SIP** | 26 5/7 weeks | 27 0/7 weeks | Not recorded |

### Expression of Wnts and other epithelial regulating molecules also vary over time and by region of intestine

Given the patterns of change of epithelial markers, we next investigated regulators of epithelial differentiation in the mouse RNASeq data. Only five members of the Wnt family were expressed at appreciable levels in the B6 mouse small intestine: *Wnt2b*, *Wnt3*, *Wnt4*, *Wnt5a*, and *Wnt9b*. Duodenal and ileal expression of *Wnt2b* were similar (Fig. 6A). Interestingly, *Wnt3* expression peaked at the same time point as the ISC markers, at day 6 in the duodenum, while it rose by day 6 in the ileum and then plateaued. Interestingly, while *Wnt3a* is used in many commercial intestinal organoid preparations, it is not expressed in the mouse small intestine*. Wnt4* expression was stable over time in the ileum and increased in the duodenum after weaning. *Wnt5a*, the prototypical non-canonical Wnt, was expressed more highly in the ileum, while *Wnt9b* was predominantly expressed in the duodenum during the first two postnatal weeks.

**Figure 6.**
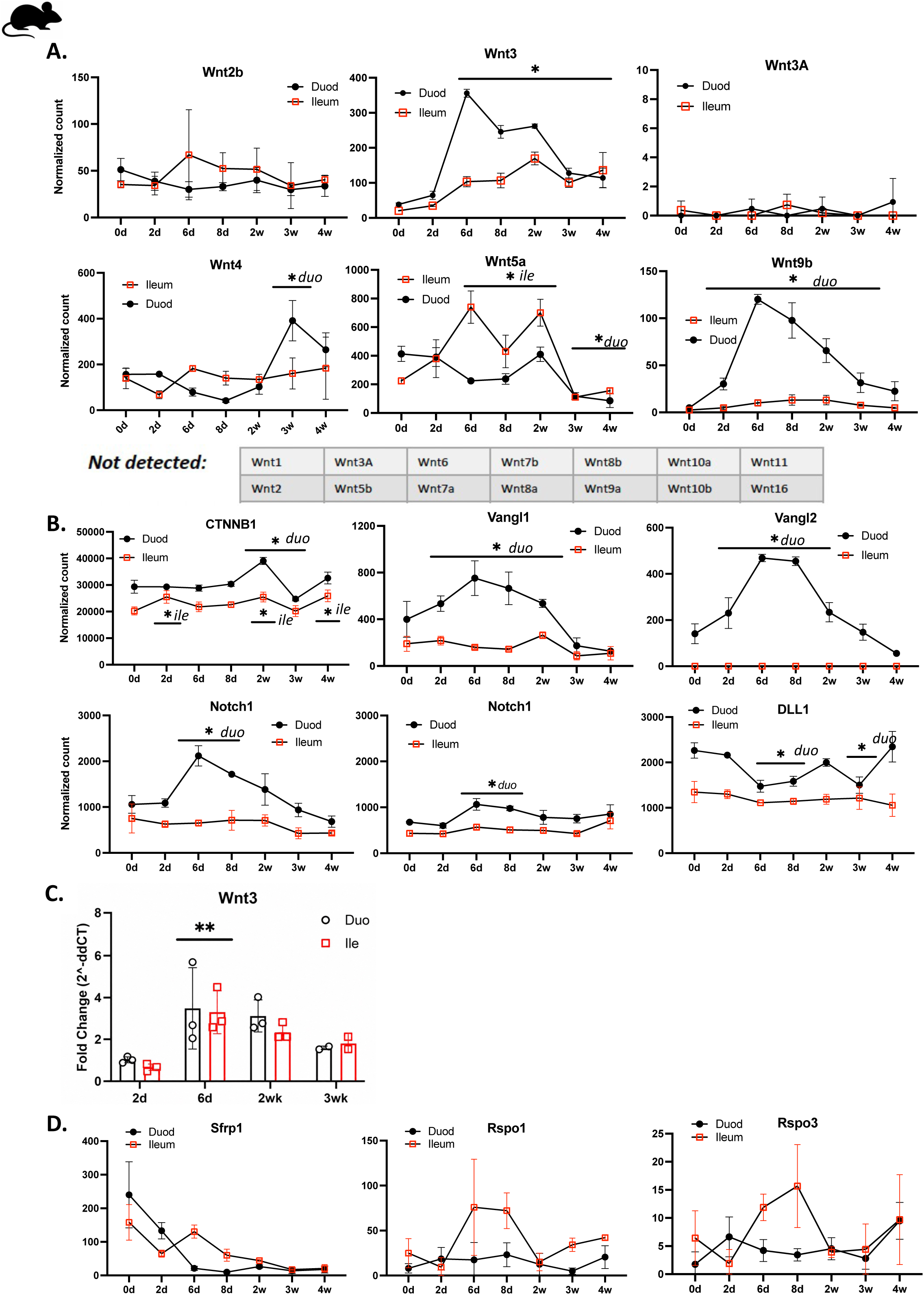
Expression of intestinal Wnts and differentiation factors. Gene expression in the duodenum (black) and ileum (red) by normalized gene count across the first month of postnatal life. Highlighted genes include Wnt family members (A) and an array of downstream regulators and trophic factors known to be important for intestinal epithelial regulation (B). Statistical analysis was by 2-way ANOVA compared to day 0. (C) qRT-PCR analysis of Wnt3 from duplicate intestinal samples at the indicated time points. *p<0.05, *duo – duodenum significant only, *ile-ileum significant only. **p<0.01

Expression of beta-catenin (*Ctnnb1*) was relatively stable over time in both segments, although it was significantly higher in the duodenum at week 2 and there was some fluctuation in the ileum (Fig. 6B). This is not altogether surprising, given that the phosphorylation state of the protein is typically the point of regulation of beta-catenin function, rather than total level of RNA or protein present (*32–34*). *Vangl1* and *Vangl2* are involved in non-canonical Wnt signaling, and both of these peaked in the duodenum at days 6 and 8 and were higher in the duodenum than ileum (in contrast to the prototypical non-canonical Wnt, *Wnt5a*, which was expressed at higher levels in the ileum). *Vangl2* was not expressed in the ileum. Expression of *Notch1 and Notch2*, which function in lineage specification, peaked in the duodenum at day 6 but remained stable in the ileum. *Dll1* (Delta-like 1), a Notch ligand, was also stable in the ileum but was significantly lower at day 6, day 8, and week 3 in the duodenum.

We again assessed select genes using qRT-PCR to assess reproducibility and consistency between assay types (Fig. 6C). *Wnt3* expression was indeed highest at day 6, but, similar to what we observed for *Olfm4* (Fig. 3C), with similar trends in the duodenum and ileum. Protein-based confirmation was not performed because Wnts are palmitoylated and antibody-based assays are unreliable.

Finally, we also evaluated genes that are associated with prenatal (E16.5) small intestinal zonation (*35*), *Wnt4* and *Srfp1* for the proximal intestine and *Rspo1* and *Rspo3* for the distal intestine (Fig. 6D), and postnatal expression was not similar to published prenatal zonation trends for these gene groups.

### Immune genes are up regulated in the mature duodenum and ileum compared to the perinatal intestine

Given that cluster 3 was highly enriched for immune-related genes, we more closely evaluated markers of the immune system. When we compared the genes most differentially expressed in the week 4 samples compared to the day 0 samples, immune genes were enriched among the significantly increased genes in both the duodenum and ileum (Fig. 7A). Several defensins and *CD19* were among the increased genes in both tissues. *Lyz1* expression was also significantly increased in the ileum. In the duodenum, insulin signaling-related genes (*Igf2, Igf2bp1*) decreased, while oxygen responsive genes also decreased in both tissues (*Hif3a*).

**Figure 7.**
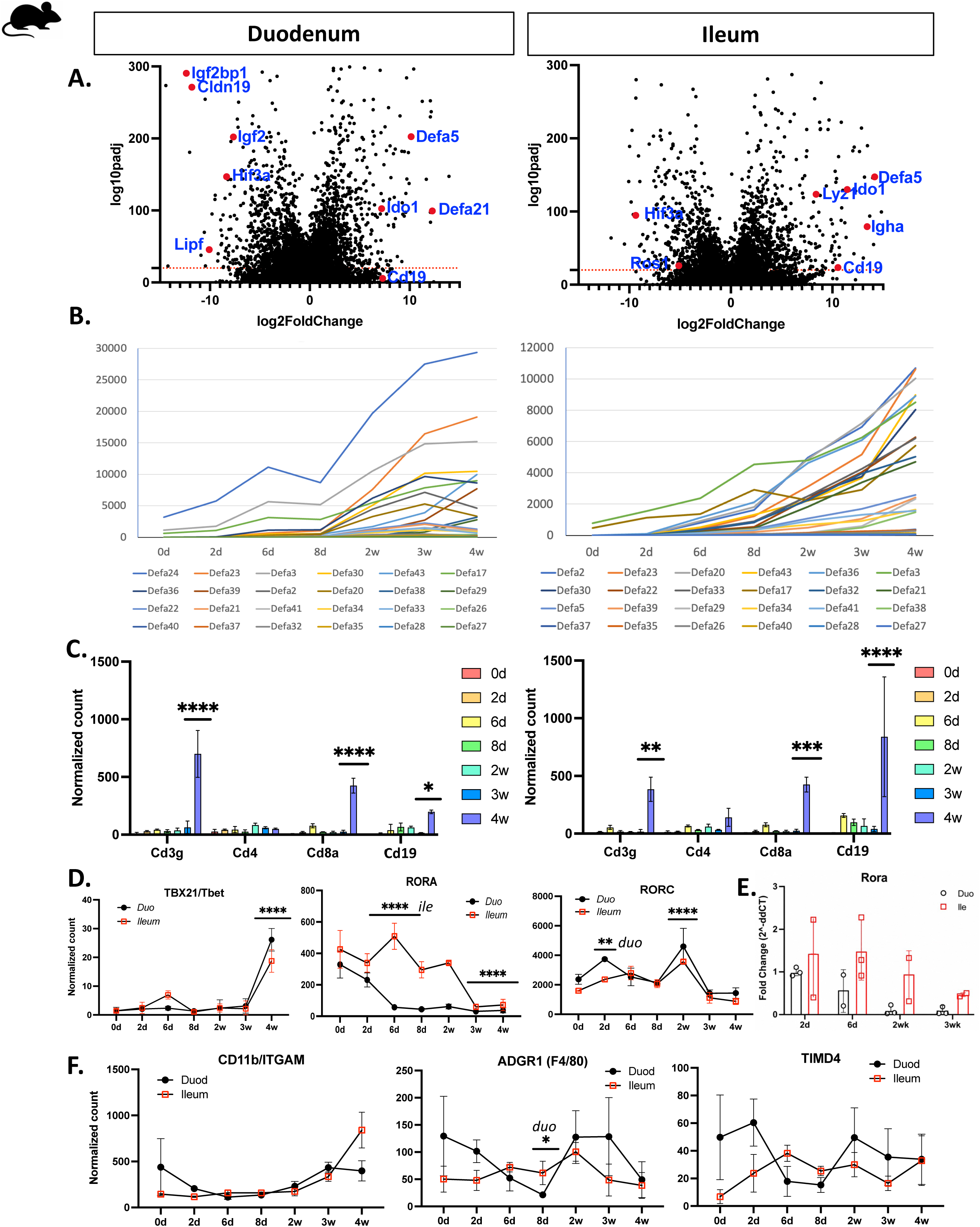
Expression of immune genes. (A) Volcano plot of all differentially expressed genes from day 0 to week 4 in the duodenum and ileum. Highlighted genes are enriched for immune regulators. (B) Gene expression of all alpha defensins in the duodenum and ileum by normalized gene count across the first month of postnatal life. Expression of adaptive immune cell markers (C) and innate lymphoid cell markers (D) by normalized gene count. (E) qRT-PCR analysis of Rora from duplicate intestinal samples at the indicated time points. (F) Expression of macrophage markers by normalized gene count. We used one-way ANOVA within each group and indicated significant values within the group as compared to day 0. *p<0.05; **p<0.01; ***p<0.001; ****p<0.0001; *duo – duodenum only; *ile – ileum only

### RNA expression of antimicrobial defensins and adaptive immune cells increase over time

Cluster 17 and the volcano plots comparing expression across all time points were highly enriched for defensins, so we also looked at these in isolation. We first compared expression of alpha defensins over time, and all defensins expressed in the duodenum and ileum increased over time (Fig. 7B, Table 1). Overall expression of defensins was higher in the duodenum than the ileum, up to 3-fold higher for some of the genes. At week 4, *Defa24, Defa3*, and *Defa23* were the highest expressed in the duodenum, whereas *Defa2, Defa23*, and *Defa 20* were the highest in the ileum. We also evaluated markers for adaptive immune cell populations (Fig. 7C). *CD3g* (T cells), and *CD8* (cytotoxic T cells) all remained low until 4 weeks. *CD4* (Th cells, intestinal macrophages) demonstrated low expression throughout the first postnatal month. *CD19* (B cells) also increased significantly at 4 weeks. Expression of T cell and B cell markers was similar between duodenum and ileum.

Expression of key gene markers for innate lymphoid cell (ILC) populations also changed over the postnatal time course (Fig. 7D). *Tbx21/Tbet* remained low until 4 weeks postnatal, suggesting that ILC1 do not populate the small intestine until later in postnatal life. *Rora*, a marker for ILC2, was higher in the immediate perinatal period and decreased over time. However, while expression fell by postnatal day 6 in the duodenum, levels remained higher in the ileum through week 2. *Rorc*, a marker for ILC3, remained more stable across development, but there was an increase in both duodenum and small intestine at 2 weeks postnatal. *Tbx21* is also expressed in inflammatory T cell populations cells, and *Rorc* in pro-fibrotic, IL-17 producing T cells, so some of the increase in *Rorc* expression may also be coming from T cells. Studies on ILC development in the postnatal intestine have been limited. We used qRT-PCR to confirm the expression pattern we saw for *Rora*, and indeed the ileum expression remained high until 3 weeks postnatal, while the duodenal expression began to decrease by 1 week postnatal (Fig. 7E).

The intestine is a site of active macrophage engagement, both from systemic macrophages and tissue-resident macrophages, which have different ontogenies (*36, 37*). *CD11b*, a marker of innate immune cells (*38, 39*), remained fairly stable across development (Fig. 7F). Expression of macrophage-specific marker *Adgr1* (F4/80) was also relatively stable, with the exception of a dip in expression in the duodenum at day 8. Meanwhile *Tim4*, a marker of tissue-resident macrophages (*40, 41*), was also stable, suggesting little fluctuation in the size of this population across early development.

We then re-evaluated the RNASeq data set using published single-cell comparator sets that identify key subpopulation marker groups. We applied an RNASeq immune cell deconvolution (*42, 43*) approach that combines key genes expressed by immune cell populations in order to estimate more precisely the changes in immune cell populations (Fig. 8). Indeed, we saw the same pattern of B cell and T cell expression increasing over time. Numbers of basophils also went up with maturation. Interestingly, there was a relatively high signature for eosinophils throughout the time points in both SI segments. Macrophage numbers appeared to decrease over time, while the signature for neutrophils increased with maturation. The total proportion of B and T cells was higher in the ileum, which is not altogether surprising given the presence of Peyer’s patches in that tissue, but outside of that difference patterns were quite similar between the two segments.

**Figure 8.**
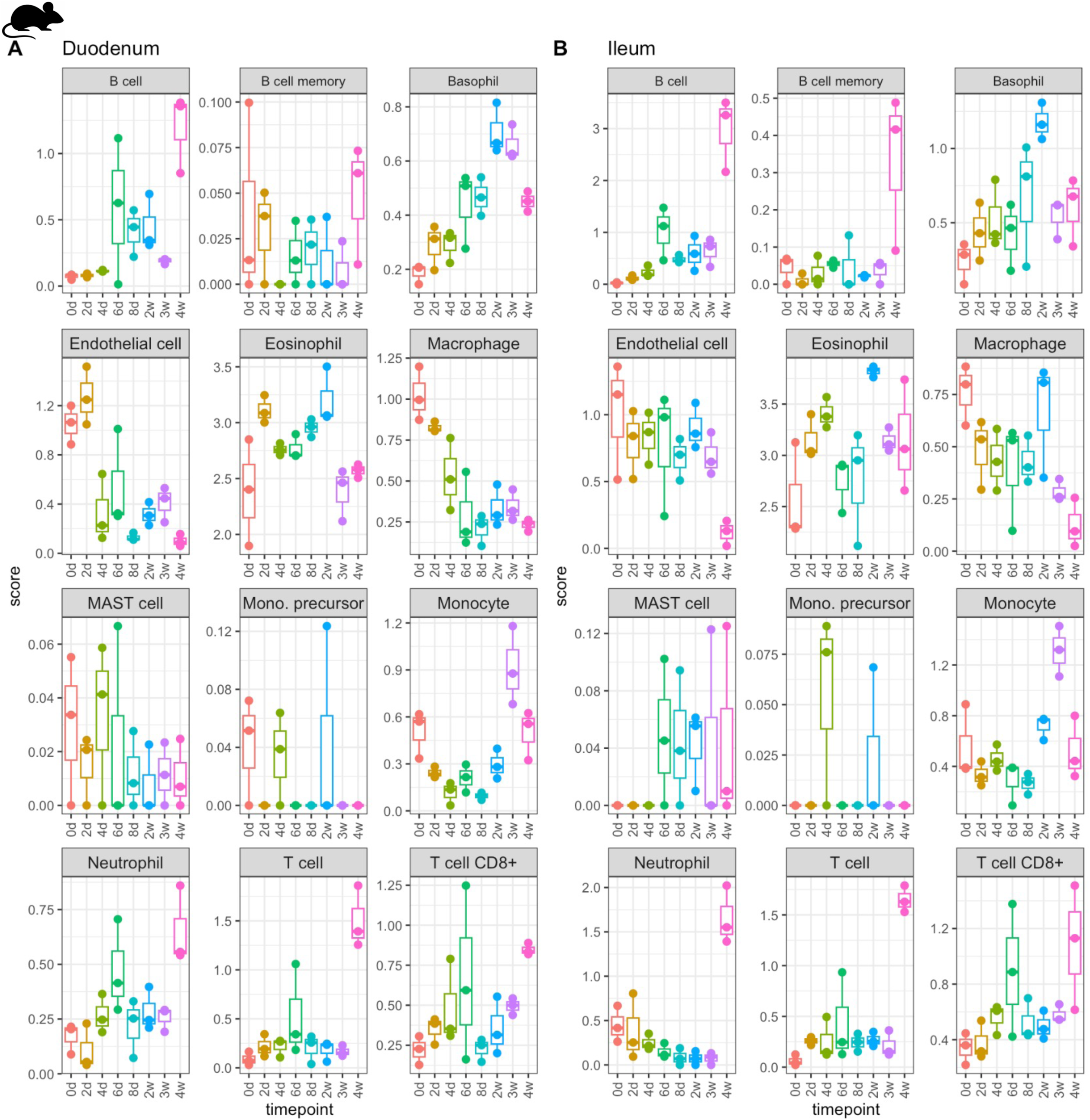
Immune cell deconvolution. Estimation of immune cell populations based on bulk RNA-seq marker gene expression in (A) duodenum and (B) ileum. The score is in arbitrary units that are comparable between samples but not between cell-types.

### Immunofluorescence analyses confirm and contrast our RNA findings

To evaluate whether the RNA changes we saw were consistent with protein changes in the developing mouse, we performed immunofluorescence (IF) of several key markers in the developing ileum. *Lyz1* expression, a marker of Paneth cell development, was low until week 3 (Fig. 3B), and IF confirmed that the protein production matched this (Fig. 9, column 1, Sup. Fig. 4A), as others have described previously (*44*). Interestingly, OLFM4 expression, a marker of intestinal stem cells, (Fig. 9, column 2; Supp. Fig. 4B) also increased steadily over time in the ileum, which was different than the RNA expression pattern, where peak *Olfm4* was seen at day 8 (Fig 3C). It was more similar to the human qRT-PCR data, which showed that organoids from older infants had higher *Olfm4* expression. We also examined the duodenum by IF and found the same pattern as in the ileum, where protein expression increased over time and was maximal at week 4 (not shown) while the RNA expression was much higher at day 6.

**Figure 9.**
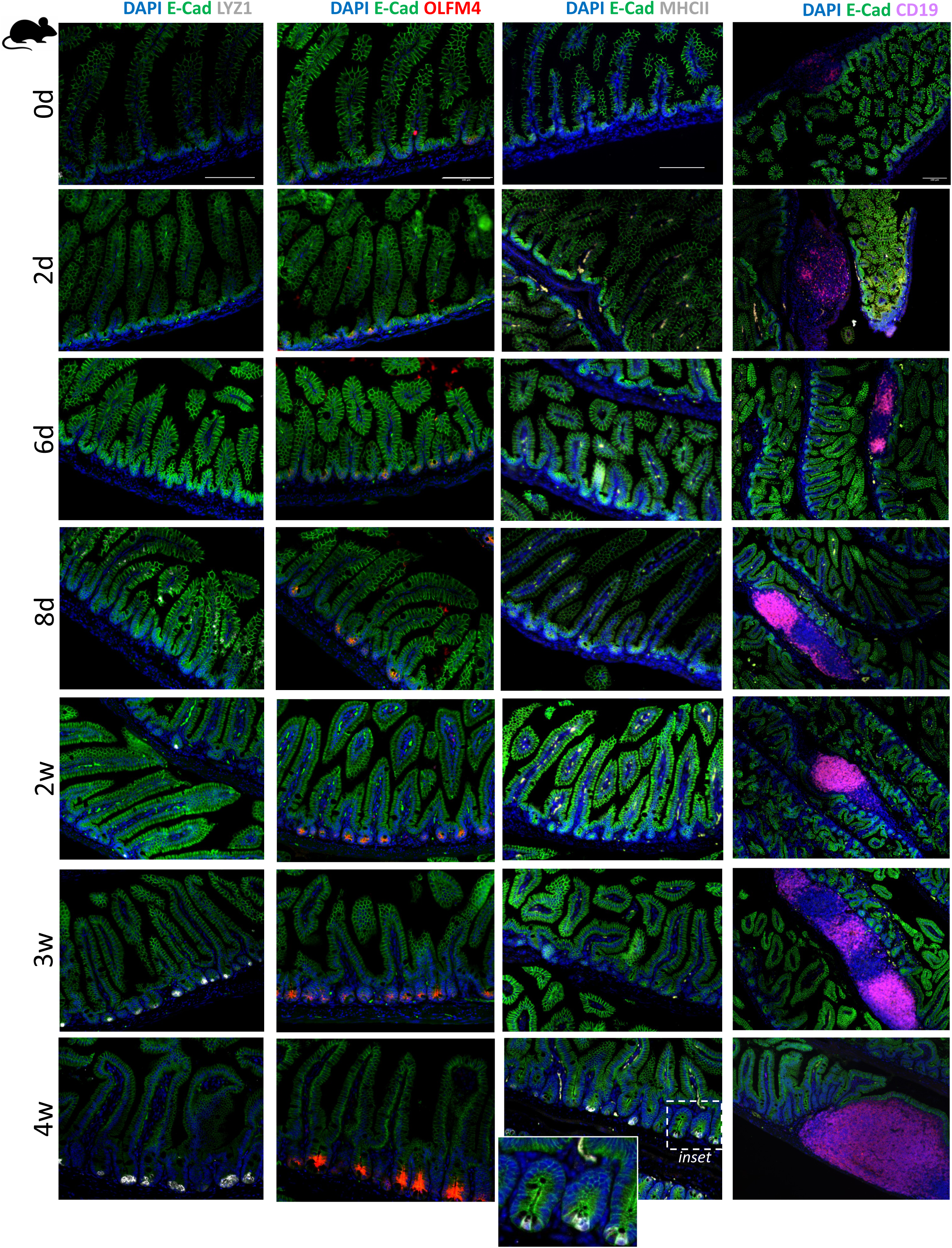
Protein expression of key epithelial and immune markers in the ileum using immunofluorescence. Ileal tissue was stained with E-cadherin (green) and DAPI (blue) as well as Lyz1 (grey, column 1), Olfm4 (red, column 2), MHCII (grey, column 3), and CD19 (magenta, column 4) at the indicated time points. Scale bar = 100 microns.

Recent advances have shown a critical role for intestinal epithelial MHCII expression in several disease processes (*45–48*). Indeed, MHCII positive cells did not emerge until week 4, when MHCII was highly expressed by certain crypt cells, likely to be ISCs based on prior reports and overlap with the OLFM4 distribution (*45*). Finally, to analyze the production of CD19, which was expressed in early time points but increased significantly at week 4, we stained tissue sections for CD19 (Fig. 9, column 4). Indeed, Peyer’s patches were small in area at day 0 and consistently enlarged over time, as described (*49*). Interestingly, early postnatal Peyer’s patch (PP) tissue was highly cellular with DAPI+ cells and few CD19+ cells, and the proportion of CD19+ increased over time as a percentage of the total PP lesions, (Supp. Fig. 3C). Postnatal development of PPs is not well described (*50*).

We then assessed expression of mesenchymal markers, which are also poorly characterized in this time period (Fig. 10). Expression of CD34, which is highly expressed in mesenchymal populations that closely associate with the ISC niche (*51*), was expressed most at the epithelium-mesenchyme interface throughout the villus, and spread as the villi grew. Podoplanin, which is a pan-fibroblast marker, was expressed highly in the submesenchymal space in a linear fashion initially, and this expression decreased with time as intravillus expression greatly increased. On the composite image, the CD34^+^ expressing cells layered on the periphery at the epithelial border and the podoplanin^+^ cells were more interior.

**Figure 10.**
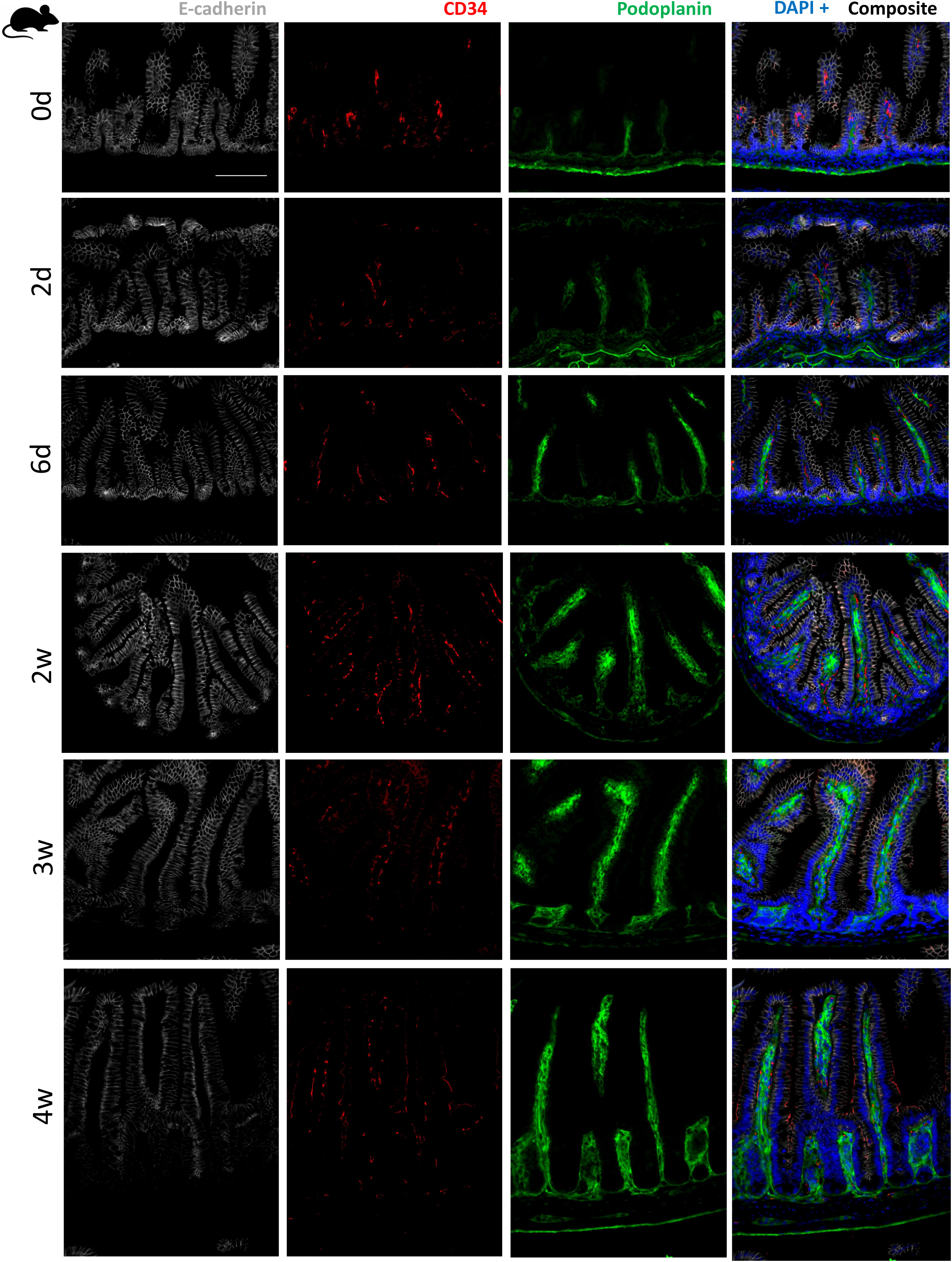
Protein expression of key mesenchymal markers in the ileum using immunofluorescence. Ileal tissue was stained with E-cadherin (gray, column 1) and DAPI (blue, in composite) as well as CD34 (red, column 1) and Podoplanin (green, column 3), at the indicated time points. Scale bar = 100 microns.

### The biological processes associated with genes differentially expressed in duodenum versus ileum changed over time

We next evaluated changes in gene expression patterns between the duodenum and ileum. Comparing genes differentially expressed in the duodenum versus ileum at all time points showed a predominance of genes related to metabolism in at day 2, day 6, 2 weeks, and 3 weeks (Supp. Fig. 3; Supp. File 4). The exception was at day 6, where there was an increase in genes related to cell proliferation, cell structures (tubules, subunits), and phosphorylation compared to the ileum. At 4 weeks, the differentially expressed genes shifted to show a predominance of immune-related gene families in the duodenum compared to ileum (Supp. Fig. 3).

## Discussion

In this study we generated an RNA catalogue of the full-thickness duodenum and ileum in the postnatal intestine of mice and analyzed gene expression patterns over time. Our study is unique in its inclusion of many time points throughout the first month of life. Previously published studies on postnatal intestinal development used targeted qPCR and IF, featured fewer time points, and focused on select functions or groups of genes (*52–54*). We also separately analyzed duodenum and ileum, and highlighted that some genes follow a similar pattern in proximal and distal small intestine, while others have distinct RNA expression patterns between the regions.

Our lab is interested in ISCs and epithelial homeostasis, particularly the role of WNT2B in control of epithelial development and homeostasis. Indeed, there were important differences in epithelial marker genes and regulators of epithelial maturation across this time course. The development of epithelial and mesenchymal lineages and the regulation of intestinal stem cells in the postnatal intestine are not well-studied. Our data demonstrate expression patterns of key postnatal cell markers, and indicate an interesting peak in ISC gene expression around postnatal day 6-8. The significance of this is currently unclear, but could potentially be a cause or result of increased crypt fission around this period as the number of structures expands. *Fgfbp1*, a marker of a reserve stem cell population(*55, 56*), in contrast, increased steadily with maturation. Epithelial deconvolution approaches showed statis expression Tuft, Goblet, and EEC cells, while Paneth cells and mature enterocytes increased with time (Fig. 4A).

Mesenchymal populations are most well studied in the context of inflammatory bowel disease and adult organisms, and are very understudied in early development. Our data, based on single gene expression (Fig. 3E), cellular deconvolution (Fig. 4B), and IF (Fig. 10) support distinct changes in fibroblast subpopulations during this critical period, where certain genes and cell types (*Pdgfra, CD34, Foxl1*) start high and then decrease as the cells spread and the mesenchyme matures. Vimentin and podoplanin RNAs are relatively stably expressed (Figs.3E, 10), while CD81 remains stable in the ileum but increases steadily in the duodenum (Fig. 3E) *Pdgfra* and *Foxl1* follow the same pattern as deconvolution data population lamina propria fibroblasts (Figs. 3E, 4B), while the subepithelial telocytes and submucosal fibroblast populations had similar expression trends during postnatal life. These populations require a lot more study in the future to understand how they overlap and differ in the perinatal period by both signature and function.

We have previously shown that WNT2B is required for normal numbers of ISC (*57, 58*), and that WNT2B deficiency leads to increased inflammatory cytokine production in the colon (*59*). In this dataset, *Wnt3* expression was substantially higher than *Wnt2b* expression in both the duodenum and ileum. *Wnt3* and *Wnt4* were the most highly expressed Wnts in the duodenum, with *Wnt4A* increasing after weaning, and *Wnt5A* was most highly expressed in the ileum (Fig. 6A). This highlights the need to consider zonation when considering the roles of trophic factors on epithelial maintenance. We also considered the markers associated with early zonation (Fig. 6A, 6D) for duodenal (*Wnt4, Sfrp1*) and ileal (*Rspo1, Rspo3*) described by a study considering late prenatal zonation in mice (E16.5 intestine) (*35*). We did not see the same pattern at day 0 and day 2, although the genes associated with ileum prenatally (*Rspo1/Rspo3*) were also increased versus the duodenum on day 6 and day 8.

Due to a high degree of homology and redundant functions, *Wnt3* and *Wnt3a* are similar in intestinal function, such that most intestinal organoid medias contain *Wnt3a*. However, our data show that *Wnt3a* is not expressed in the mouse SI, at least through 1 month of age (Fig. 6A). These Wnts are highly homologous and share some functional redundancy (*60–62*), however, their nucleotide sequences are only 55% identical with large regions of dissimilarity and whether they have any functional differences in controlling intestinal homeostasis is yet to be interrogated. Researchers who use organoids should be aware of this nuance.

The immune system is rapidly developing in the postnatal period, as the intestine first encounters the microbiome and is exposed to both microbial and food-related antigens. Despite initiation of milk feeding within hours after birth, adaptive immune cell markers did not robustly increase until week 4 (Fig. 7C, Fig. 8). Whether this is related to the introduction of standard rodent chow at day 21 in our mice or the resultant decrease in breastmilk consumption, or something else entirely, cannot be determined from our approach. Intestinal macrophage markers remained static across the first postnatal month (Fig. 7F), which may be due to the predominance of tissue-derived macrophages in this organ (*40, 63*). The changes in expression of markers specific for ILC populations were interesting. Not unexpectedly, the ILC1 markers were expressed on a much lower level than ILC2 or ILC3 markers (Fig. 7D, 7E). Expression of ILC2 marker *Rora* was higher in early time points, and expression was more persistent in the ileum compared to the duodenum. No dedicated gene marker has been identified to date for ILC1 and ILC3, however, so expression of these markers may be confounded by T cell gene expression. We were unable to analyze ILC subsets in the immune deconvolution data.

Peyer’s patch size increased over time, as previously described, but interestingly the proportion of B cells within the Peyer’s patches (PPs) relative to the total area increased significantly over time, suggesting rapid growth of the germinal centers in the postnatal period (Fig. 9, Supp. Fig. 4). Others have shown that the postnatal microbiome drives development of PPs, and that early postnatal antibiotic exposure decreases the number of intestinal B cells in flow cytometric analyses, however they only examined samples from P20 onward. The effects of antibiotics, microbial composition, and other postnatal exposures on expansion of GCs requires additional study.

The main limitation of this study is that most data were generated by bulk RNA-Seq. We have done some select confirmatory studies using qRT-PCR and immunofluorescence but have not recapitulated the majority of the dataset by other means. This dataset is intended as a starting point for our lab and others when considering novel hypotheses, and secondary approaches should always be used to further interrogate the RNA expression in a given experimental system. Furthermore, total mRNA does not equate to transcriptionally active mRNA or to protein levels, so other assays would be needed to remark on those aspects of translational control. Another caveat is that we intended to collect data from day 4, but these samples were not of high enough quality, so we omitted that data. Some dynamic changes occur for ISC genes around day 6, so it is not yet clear whether these markers change soon after day 2 or closer to day 6. Finally, this study is done in C57Bl/6 mice, so applicability to other backgrounds cannot be assumed.

It is important to keep in mind that these samples are all under homeostatic conditions, and the absence or abundance of a gene in this dataset may not be reflective of its expression during stress or injury conditions. Furthermore, exposures are changing over the period we queried, so microbial changes and nutrient changes are occurring but should be relatively consistent in all postnatal mice. We were curious about the natural sequence of events in the normal postnatal environment under routine conditions, so we did not manipulate any of these exposures. Of course, the microbiome is well known to play an important role in development of the epithelial transcriptome and in postnatal methylation, as other groups have demonstrated (*64*).

While technological advances have allowed the advent of single-cell sequencing approaches for understanding mouse transcriptomics, isolation and representative preparation of both intestinal epithelium and mesenchyme simultaneously remains a challenge, which can make comparing between segments convoluted. Furthermore, rare cell types and RNAs present at a lower frequency can be difficult to capture in single-cell sequencing (*65*). This study serves as a launch pad for future investigations, and both single-cell and spatial analyses in humans and mice would be additionally informative for the field.

In sum, the first postnatal month of life is one of the most dynamic times for intestinal growth and crypt expansion. Characterizing the normal developmental sequence can help us to identify developmental aberrations in neonatal intestinal diseases that may drive pathogenesis. We hope this dataset can serve as a reference point for others studying specific genes or biological processes during postnatal health and disease.

## Methods

### Research Ethics

The study was approved by Boston Children’s Hospital Institutional Review Board for the human subjects research (BCH IRB 10-02-0053 and P00027983) in accordance with the Declaration of Helsinki. All animal experiments were done in accordance with Boston Children’s Hospital Institutional Animal Care and Use Committee (IACUC).

### Sex as a biological variable

The role of sex was not considered as a biological variable in this study, and male and female samples were used indiscriminately for all analyses. This was mainly owing to the high cost of RNA-Seq and the difficulty in telling male from female mice at the earliest time points of the study. We do provide data on sex of the human samples, when able.

### Reagents

A table of reagents used in this study, including antibodies and qPCR primers, is included in Supplementary Table 1.

### Mouse model

C57/Bl6 mice (Jackson labs) mice were used for experiments, and for RNA-Seq experiments, we used large litters and collected mice across postnatal ages to the extent possible (usually 2-3 time points per litter), to limit potential variation in microbial exposures. Mice were housed in standard germ-free conditions and kept with the dam until 21 days postnatal and then transitioned to standard rodent chow. We used both male and female mice for all experiments.

### Time course sample collection

Mice were euthanized at the following time points: day 0, day 2, day 4, day 6, day 8, 2 weeks, 3 weeks, and 4 weeks. For the purposes of RNA-Seq, 2cm of proximal intestine immediately after the pylorus was collected for the “duodenum” sample and 2cm of distal intestine immediately before the ileocecal junction was collected for the “ileum” sample. For the histology samples, we isolated the full small intestine and then transected at the mid-way point to create proximal (duodenal-jejunal, called “duodenum” throughout) and distal (“ileum”) intestinal segments. The intestine was cleaned of feculent material using cold PBS and 4% paraformaldehyde (PFA), and then tissue was opened longitudinally, rolled, and placed in a fixation cassette (Swiss roll method). All samples were Swiss-rolled with orientation of the rolls in a proximal to distal fashion, in order to be able to compare consistently between segments. We analyzed only the segments that corresponded to the duodenum and ileum for the histology, in order to maintain consistency with the RNA data.

### RNA-Seq

Intestinal tissue was placed into Trizol LS (ThermoFisher) and homogenized with a bead homogenizer. RNA was isolated using the Direct-zol kit (Zymo Research) according to the manufacturer’s instructions. The library was prepared by the Dana Farber Next Generation Sequencing Core Facility using a KAPA library quantification kit (Roche). Sequencing was done on an Illumina NovaSeq 6000 with total RNA.

All samples were processed using a bulk RNA-seq pipeline implemented in the bcbio-nextgen project (bcbio-nextgen,1.2.8-1c563f1). Gene expression was quantified using Salmon (version 0.14.2) (*66*) using the mm10 transcriptome (Ensembl). Differentially expressed (DE) genes between the duodenum and ileum at specific time points were identified using DESeq2 (version 1.30.1) (*67*). The DE threshold was set at adjusted P-value <= 0.005. All analyses performed in R utilized R version 4.0.3.

### Analysis of Gene Expression Families Over Time

We analyzed gene expression over time on the duodenum and ileum samples separately using the LRT method in DEseq2. After identifying differentially expressed (DE) genes, we the degPatterns() function from the DEGreport package (version 1.34.0, citation – Pantano L (2024). *DEGreport: Report of DEG analysis*. R package version 1.42.0, http://lpantano.github.io/DEGreport/.) to identify “clusters” of genes with similar expression patterns across the time points in each tissue using the top 3000 DE genes. We visually analyzed the expression patterns to identify gene groups that fluctuated from the time of NEC susceptibility (less than 2week of age) to the time when mice become resistant to NEC (2 weeks of age onward). We performed functional analysis using the R package clusterProfiler (version 3.18.1) (*68, 69*) on these gene sets to identify GO biological processes or KEGG pathways that are enriched in each cluster of genes. We then expanded the gene list to include other DE genes with the same pattern of expression of the groups we identified using DEGreport (version 1.34.0sak, citation – Pantano L (2024). *DEGreport: Report of DEG analysis*. R package version 1.42.0, http://lpantano.github.io/DEGreport/.)), and performed enrichment analysis for GO terms (biological process).

### Immune Deconvolution

Immune cell deconvolution was performed using the mMCPcounter method (version 1.1.0)(*30*) as implemented in the immunedeconv R package (version 2.1.0)(*70*) using R version 4.4.0. In brief, gene expression counts were normalized to transcripts per million (TPM) and used as input to the mMCPcounter function, which quantifies cell populations based on transcriptomic markers. This function provides cell-type scores for each sample in arbitrary units that are comparable between samples but not between cell-types.

### Epithelial and Mesenchymal Deconvolution

Deconvolution analyses were performed using the transcripts per million (TPM) normalized gene expression matrix and a list of marker genes as input to the mMCPcounter tool version 1.1.0 (*30*) using R version 4.5.1. This reference dataset was also used for the immune deconvolution. Epithelial cell markers were taken from Haber et al., 2018, Supplemental Table 4, a single cell dataset from adult mouse epithelium as the reference library (*71*). Mesenchymal cell markers were taken from Johnson et al., 2025 preprint, Figure 3 analyzing single cell transcripts from the whole length of the small intestine of late embryonic stage mouse intestine at P16.5 (*29*). mMCPcounter returns scores in arbitrary units that are comparable between samples but not between celltypes.

### Pathway overrepresentation analyses for duodenum vs ileum

Differentially expressed gene lists of duodenal results relative to ileum at various time points were uploaded to WebGestalt 2019 (*72*). Selected parameters were overrepresentation analysis, KEGG pathway, and Illumina mouse (*13*).

### Biopsy Samples from Human Infants

Biopsied tissue from endoscopic resections or surgeries was fixed in formalin and embedded in paraffin. All samples were collected solely for clinical purposes, and we used them in a retrospective fashion. Unstained slides were prepared by the clinical histology core at Boston Children’s Hospital.

### Establishment of Human organoids

Human intestinal organoids were generated using StemCell® protocol that was adapted as follows. Tissue vials containing about 10 biopsy sized pieces of small intestine that were cryopreserved in 10% DMSO in Fetal Bovine Serum (FBS) were quickly thawed in a water bath. The thawed tissue was transferred into a 15 ml conical containing 5 ml of ice-cold Phosphate Buffered Saline (PBS). The conical was centrifuged at 290 rcf for 5 minutes at 4°C. The supernatant was discarded, and the wash step with PBS was repeated. After supernatant was aspirated, tissue with about 1 mL of PBS was transferred into a 1.5 mL Eppendorf under a sterile hood. Using scissors, the pieces of tissue were minced until they were about 1-2 mm in diameter. Tissue suspended in PBS was transferred to a new 15 mL conical and the PBS was aspirated once the tissue settled.

### Propagation of Human Organoids

Organoids were resurrected and expanded from frozen prior to use. Briefly, vials were thawed in a 37°C water bath until just starting to liquefy and then rinsed with Dulbecco’s modified eagle’s media/F12. They were plated in 24-well plates within 50μL of Matrigel (Corning), allowed to solidify in a 37°C incubator for 10 mins, and then 500μL of human small intestinal organoid media was added (Supplementary Table 2). Media was changed every 2-3 days and organoids were passaged roughly once a week until used for experiments. All experiments were done at passage 15 or lower.

### Quantitative PCR

The Trizol extraction method was used to purify total mRNA from duodenal enteroid samples. Quantity and quality were assessed using Nanodrop 2000 (Thermofisher Scientific, MA), then cDNA was synthesized using the High Capacity cDNA Reverse Transcription Kit (Thermofisher Scientific, MA). Quantitative RT-PCR was performed using TaqMan Universal PCR MasterMix and Taqman Gene Expression Assays (Thermofisher, MA) for the indicated genes (Supplementary Table 1).

### In-situ hybridization

RNAscope Multiplex Fluorescent Reagent Kit v 2 (Advanced Cell Diagnostics, ACD) method was used to localize mRNA expression in FFPE intestinal ileum sections according to a modified version of the manufacturer’s protocol. First, the slides were baked at 60° C for 1 hour and then washed twice with Xylene (Sigma Aldrich) and twice with 100% ethanol for five minutes each. Next, the sections were treated with hydrogen peroxide reagent for 10 minutes RT, washed twice with distilled water, and hot-rinsed with boiling distilled water for 15 seconds, followed by incubation with 1X Target Retrieval Buffer at 99°C for 5 minutes in steamer. Afterward, the slides were hot-rinsed in distilled water, dehydrated in 100% ethanol, and dried for 5 minutes RT. The sections were then incubated with Protease Plus Reagent at 40°C for 30 minutes, washed twice with 1X Wash Buffer, and hybridized with LGR5 probe at 40°C for 2 hours. After hybridization according to the manufacturer’s protocols, the sections were washed twice, incubated with DAPI (1:1000) for 15 minutes RT, washed again, and finally sealed with a coverslip. ACD designed the probe used in this study (Supplementary Table 1). Images were taken using a Zeiss LSM 980 with Airyscan microscope and processed using ImageJ Fiji software(*73*).

### Immunofluorescence

Tissues were fixed in 4% PFA overnight at 4°C, washed in 50% ethanol for 1h, and then transitioned into 70% ethanol. Tissues were embedded in paraffin and sectioned (4-6μm). Slides were prepared for staining using Trilogy buffer (Sigma Aldrich) in a pressure cooker for 15 minutes according to the manufacturer’s recommendation. Slides were then rinsed with boiling Trilogy buffer for 5 minutes, washed with deionized water, and then rinsed further in PBS. Blocking buffer was added to each slide (blocking buffer: 0.3% Tween-20, 10% normal donkey serum, 0.05% bovine serum albumin in PBS) and incubated at room temperature for 1 hour, gently shaking. Primary antibody (Supplementary Table 1) was then added in blocking buffer at 4°C overnight. Slides were then washed in PBS and incubated with the secondary antibody for 1 hour at room temperature. DAPI (1:1000) was added for 10 minutes and then the slides were washed again and sealed with a coverslip. Images were taken using a Nikon Eclipse 90i Fluorescence microscope and processed using ImageJ Fiji software (*73*).

### Statistical Analyses

Statistical analysis of RNA-Seq data is described separately above. For experiments comparing two groups (e.g., duodenum vs ileum), the Student’s t test was used in PRISM 9 (GraphPad Software). For experiments assessing expression changes over multiple time points, ANOVA was used and individual points were compared to the day 0 baseline sample. Data were assumed to be normally distributed. For all experiments, p<0.05 was considered significant.

## Support

This work was supported by the National Institutes of Health NIDDK K08DK120871 (AEO), NIH P30DK034854-36 (Harvard Digestive Disease Center Pilot award, AEO), Boston Children’s Hospital Office of Faculty Development/Basic & Clinical Translational Research Executive Committees Faculty Career Development Fellowship (AEO), and the Charles C. Hood Foundation (AEO). Work by RK and SHS was supported in part by the Harvard Stem Cell Institute.

## Data Access

Metadata, raw sequencing data, and count matrices have been uploaded to GEO, GSE281618. Processed files (DE gene lists, and searchable gene clusters) are included as supplementary files.

## Supporting information

Supplemental Table 1 reagents

Supplemental Table 2 organoid media

## Abbreviations

ANOVA: analysis of variance
DE: differentially expressed
NEC: necrotizing enterocolitis
ISC: intestinal stem cells
ILC: innate lymphoid cells
IF: immunofluorescence
PP: Peyer’s patch
PFA: paraformaldehyde
MHCII: major histocompatibility complex class II
RT: room temperature
SIP: spontaneous intestinal perforation
PCA: principal component analysis
FFPE: formalin-fixed paraffin-embedded

## Acknowledgements

We would like to thank the infants who contributed tissue samples for this study and their families. The authors would also like to thank Nicholas Makoganov for assisting with some of the technical aspects for this manuscript.

## Author contributions

Authorship was indicated by order of greatest to least contribution, with the exception of the three senior authors, who were listed last in order of least to greatest contribution.

All authors read and critically reviewed the manuscript and approved of the final manuscript. In addition:

LFO performed RNAScope experiments and assisted with data collection and manuscript preparation.

SW performed some experimental assays and data analyses.

VSD contributed to experimental assays for immunofluorescence experiments.

RSK performed the bioinformatics analysis of the RNA-seq data and assisted with drafting and editing the manuscript.

SJHS provided mentorship and bioinformatics expertise, aided with data management, and critically edited the manuscript.

AWH assisted with reviewing human infant cases and identifying appropriate pathologic specimens.

ESP performed the bioinformatics analysis of the immune deconvolution data.

OO generated fetal and human neonatal organoids.

JG provided human histology specimens and expertise.

DTB provided mentorship and topical expertise, and critically edited the manuscript.

LK provided human organoids, provided topical expertise, and critically edited the manuscript.

AEO conceptualized the project, performed experiments, and drafted, edited, and revised the manuscript.

## Data availability statement

The synthesized transcript datasets generated and analyzed during the current study are included in this published article and its Supplementary Information files. The raw sequencing data is available in GEO with accession GSE281618.

## Additional Information

The authors report no conflicts of interest or competing interests related to this study.

