## Supplemental Figures for "Cataloguing the mouse small intestinal transcriptome in duodenum and ileum during the first postnatal month"

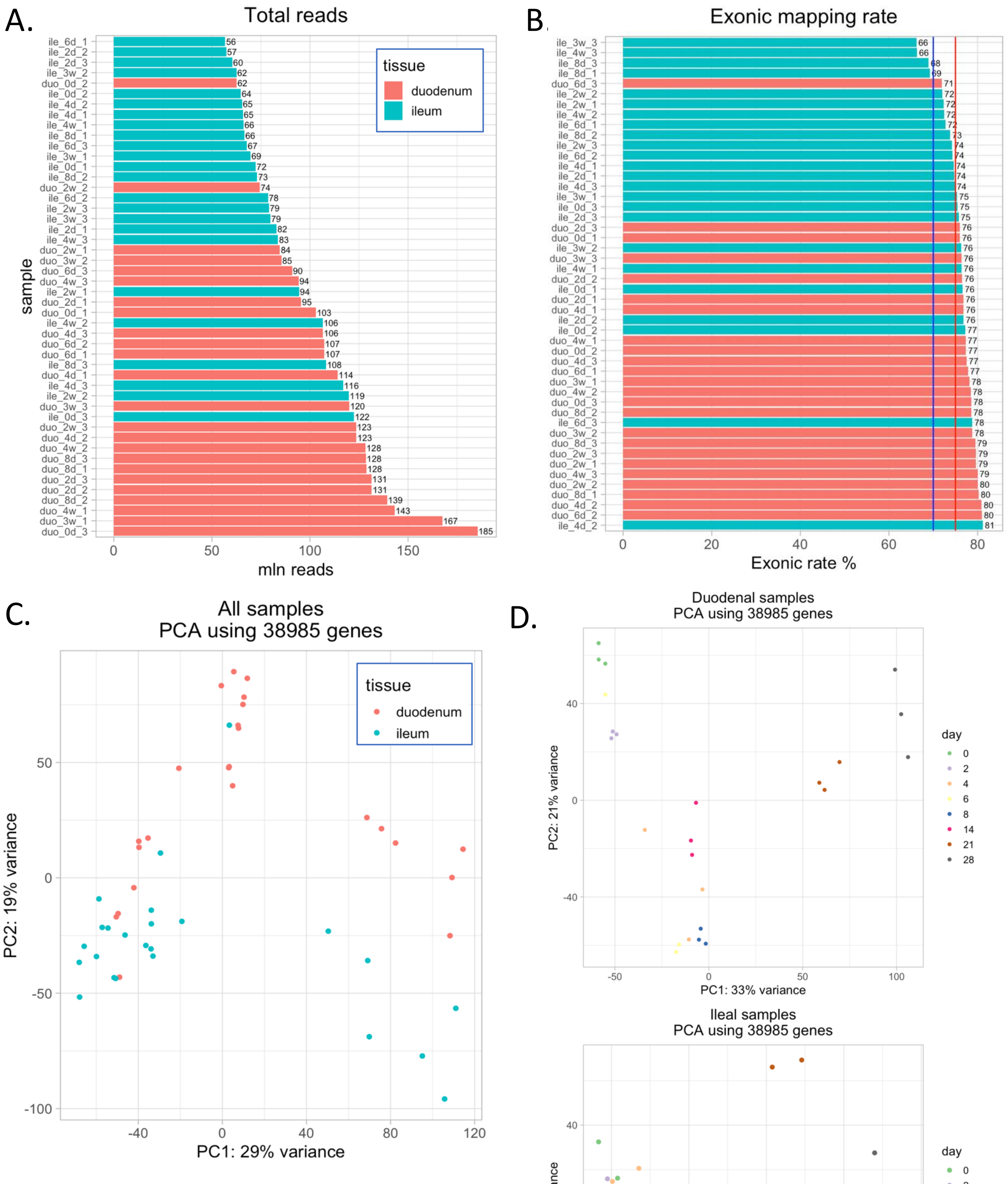

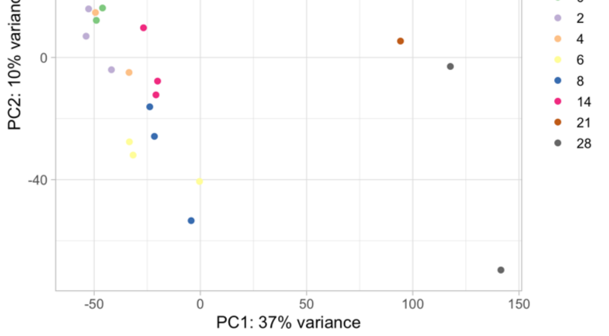


**Supplemental Fig. 1 Quality analysis of RNASeq dataset**. A. Total reads per sample, x-axis is millions of reads. B. Exonic mapping rate, color scheme is the same as for (A). C. Principal component analysis (PCA) of all samples labelled by anatomic source. (D) Clustering by timepoint for duodenal (top panel) and ileal (bottom) samples.

### Duodenum DE gene trends

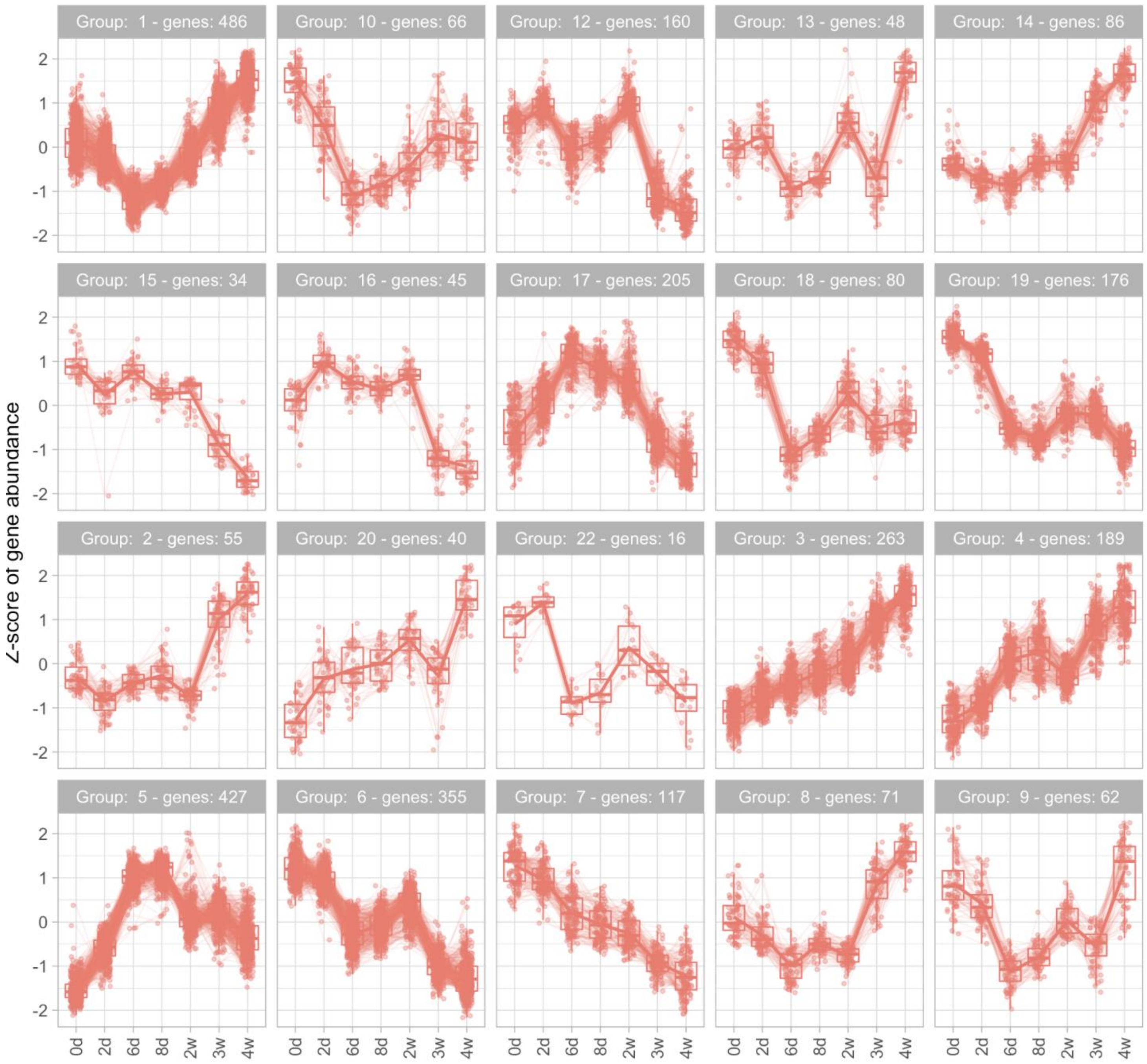


**Supplemental Fig. 2 Differentially expressed gene trends, duodenum.** We analyzed for trends in significantly different gene expression across the time points. The z-score of gene abundance is displayed, adjusted P-value threshold used <= 0.005

### Ileum DE gene trends

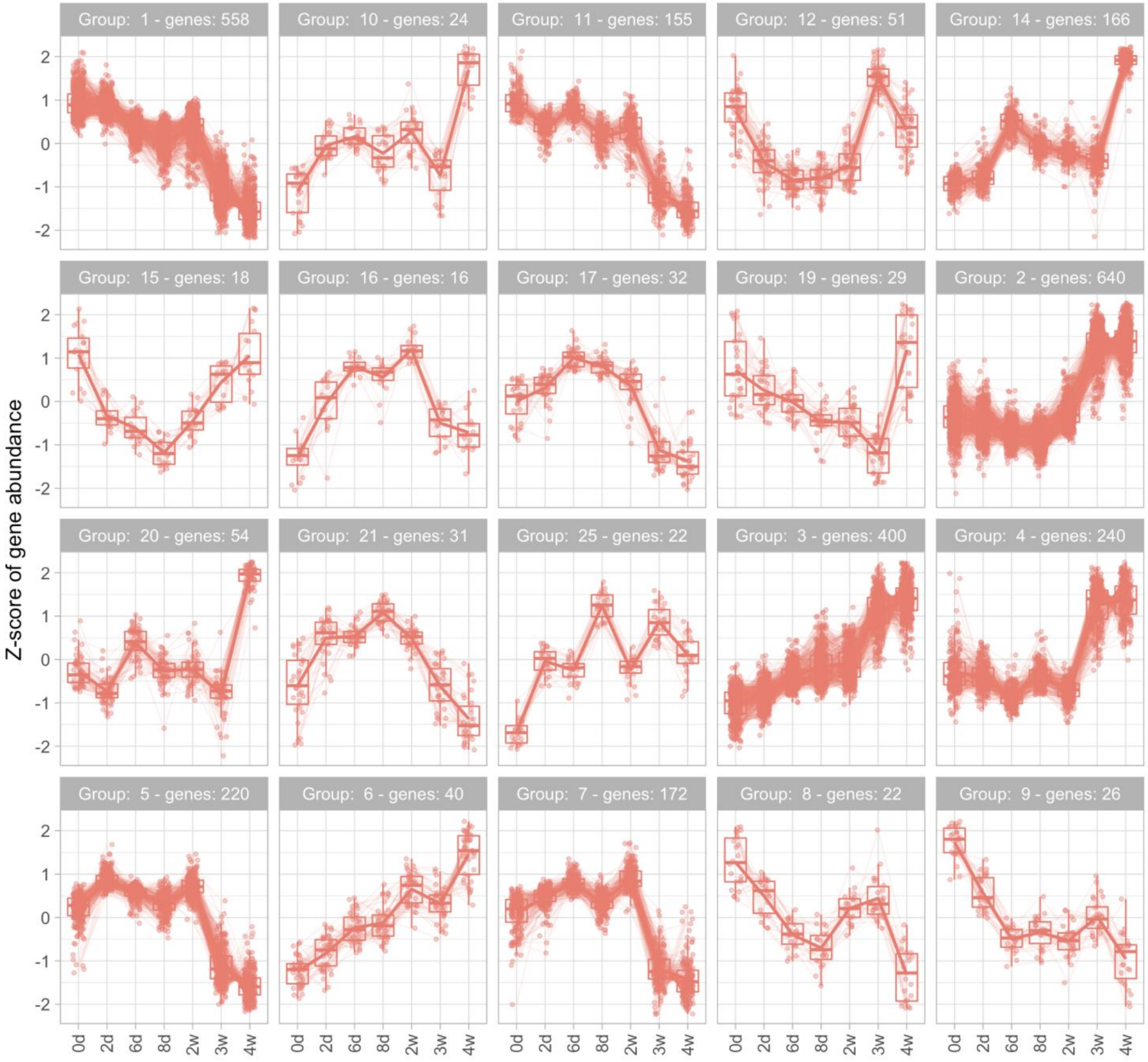


**Supplemental Fig. 3 Differentially expressed gene trends, Ileum.** We analyzed for trends in significantly different gene expression across the time points. The z-score of gene abundance is displayed, adjusted P-value threshold used <= 0.005

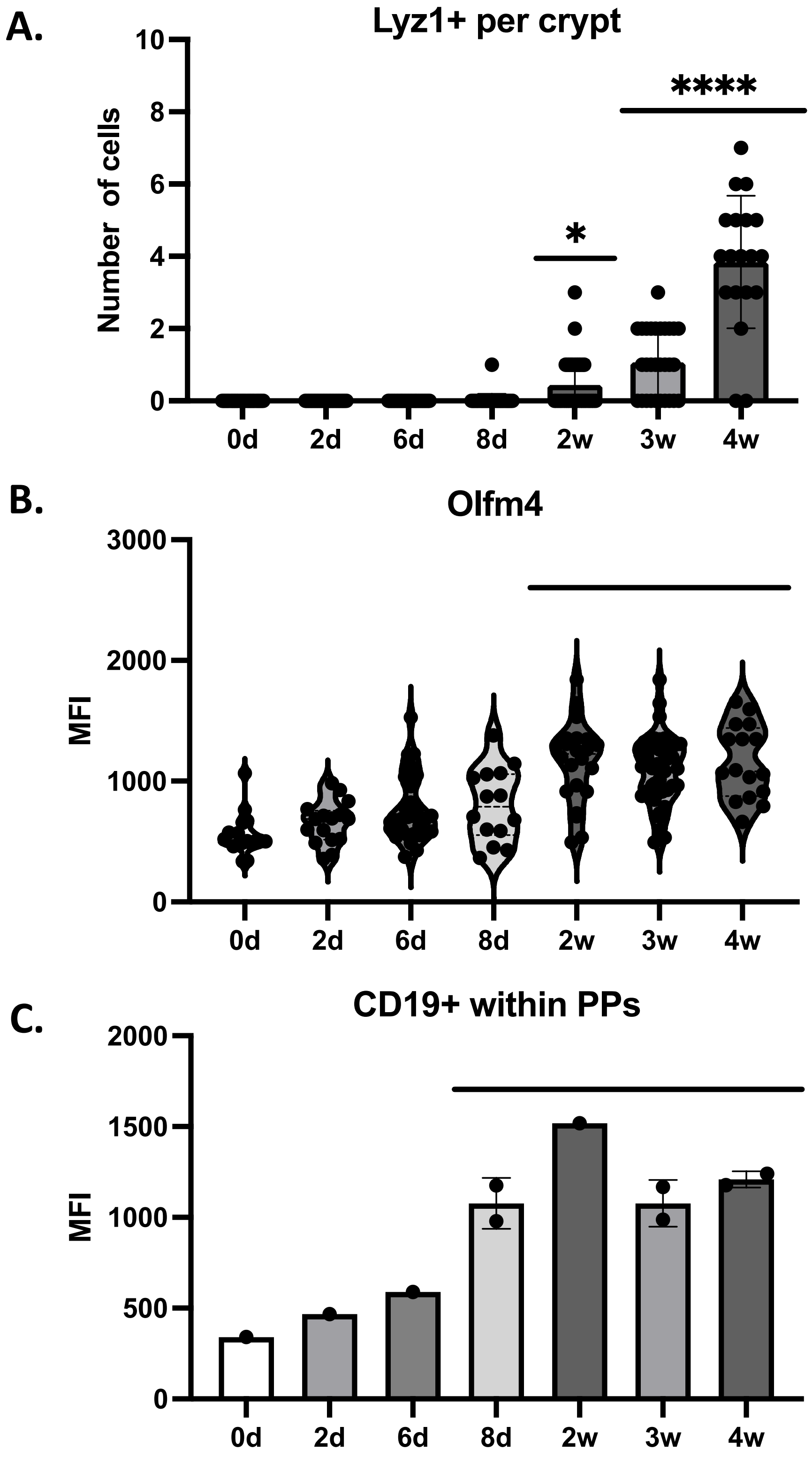


**Supplemental Fig. 4 Quantification of immunofluorescence assays** (A) Mean fluorescence intensity of Olfm4 expression in crypt bases. (B) Number of Lyz+ positive cells per crypt. (C) MFI of CD19 expression within the Peyer's patches. * $p < 0.05$ **** $p < 0.0001$

### Duodenum vs Ileum at individual time points

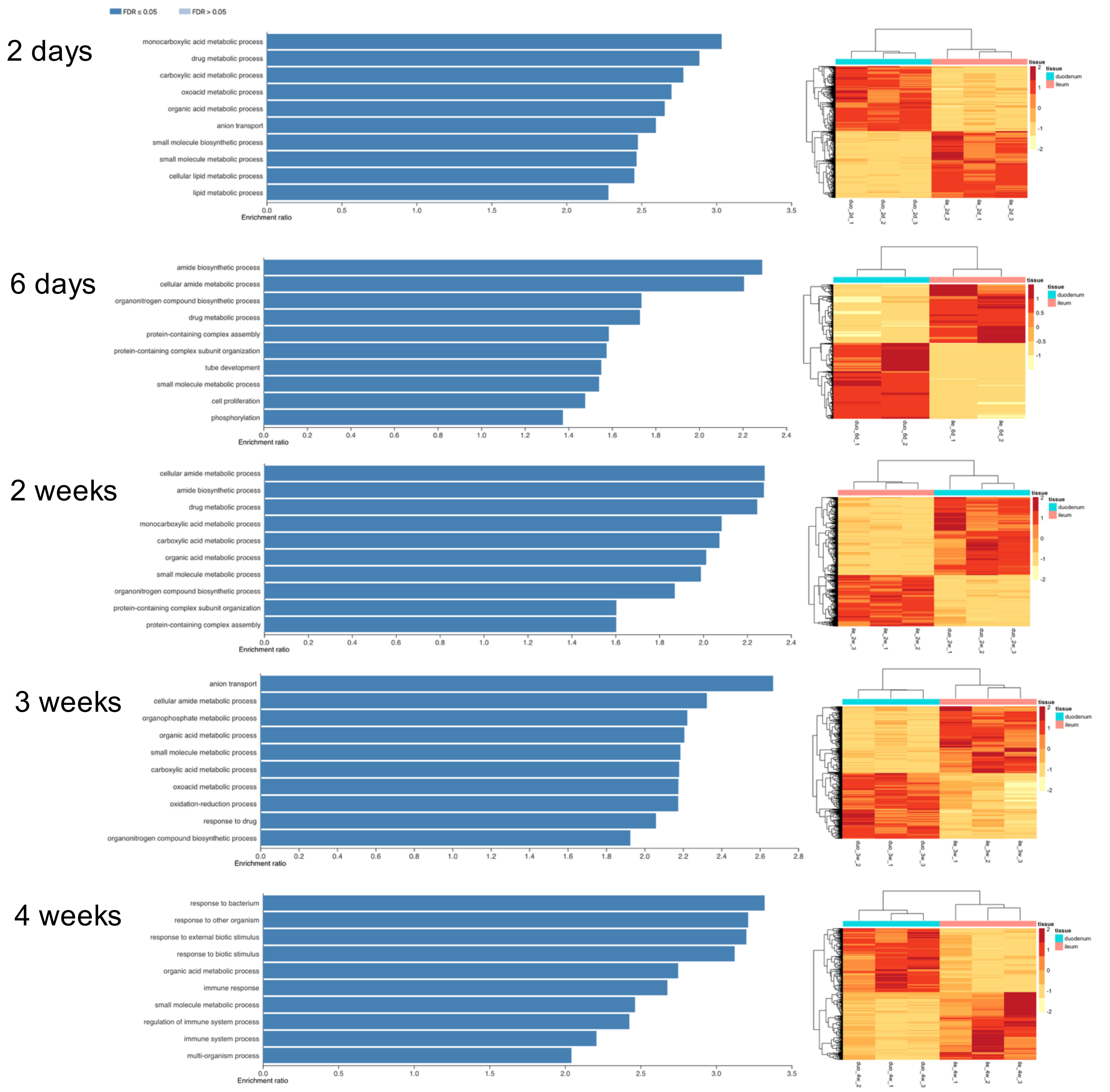


**Supplemental Fig. 5 Biological process gene overrepresentation analysis of duodenum relative to ileum.** The geneset enrichment results are displayed by timepoint.
