## Supplemental Table 2 organoid media for "Cataloguing the mouse small intestinal transcriptome in duodenum and ileum during the first postnatal month"

**Supplemental Table 2 Human Small Intestinal Organoid Culture Media**

**Human Ileum Media (100mL)**

| **Component** | **Volume** | **Catalog #** | **Final Concentration** | **Purpose** |
| --- | --- | --- | --- | --- |
| NRW Conditioned Media | 65mL | HDDC Organoid Core | 65% | Contributes Noggin, Rspondin-1, and Wnt3a for stem cell proliferation |
| Advanced DMEM/F-12 | 30mL | Life Technologies 12634028 | 30% | Contributes glucose, NEAAs, sodium pyruvate, phenol red |
| Glutamax (100X) | 1mL | Life Technologies 35050061 | 1X | Contributes L-glutamine to support growth |
| HEPES (1M) | 1mL | Life Technologies 15630080 | 10mM | pH buffer for CO2 changes |
| Primocin (50mg/mL) | 200uL | Invivogen ant-pm-1 | 100ug/mL | Antibiotic |
| Nicotinamide (1M) | 1mL | Sigma N0636 | 10mM | Vitamin A derivative |
| Normocin (50mg/mL) | 200uL | Invivogen ant-nr-1 | 100ug/mL | Antibiotic |
| B27 Supplement (50X) | 1mL | Life Technologies 12587010 | 0.5X | Growth supplement |
| N2 Supplement (100X) | 500uL | Life Technologies 17502001 | 0.5X | Growth supplement |
| N-Acetyl-Cysteine (500mM) | 100uL | Sigma A7250 | 500uM | Antioxidant |
| A-83-01 (500uM) | 100uL | Sigma SML0788 | 500nM | TGF-beta inhibitor |
| SB202190 (30mM) | 33.2uL | Sigma S7067 | 10uM | p38 MAPK inhibitor |
| EGF(500ug/mL) | 10uL | Peprotech 315-09 | 50ng/mL | Epidermal growth factor, stimulates mitosis |
| Gastrin (100uM) | 10uL | Sigma G9145 | 10nM |  |
| Prostaglandin-E2 (1mM) | 1uL | Peprotech 3632464 | 10nM |  |
